# Skeletal muscle BMAL1 is necessary for transcriptional adaptation of local and peripheral tissues in response to endurance exercise training

**DOI:** 10.1101/2023.10.13.562100

**Authors:** Mark R Viggars, Hannah E Berko, Stuart J Hesketh, Christopher A Wolff, Miguel A Gutierrez-Monreal, Ryan A Martin, Isabel G Jennings, Zhiguang Huo, Karyn A Esser

## Abstract

**Objectives:** In this investigation, we addressed the contribution of the core circadian clock factor, BMAL1, in skeletal muscle to both acute transcriptional responses to exercise and transcriptional remodelling in response to exercise training. Additionally, we adopted a systems biology approach to investigate how loss of skeletal muscle BMAL1 altered peripheral tissue homeostasis as well as exercise training adaptations in iWAT, liver, heart, and lung of male mice.

**Methods:** Combining inducible skeletal muscle specific BMAL1 knockout mice, physiological testing and standardized exercise protocols, we performed a multi-omic analysis (transcriptomics, chromatin accessibility and metabolomics) to explore loss of muscle BMAL1 on muscle and peripheral tissue responses to exercise.

**Results:** Muscle-specific BMAL1 knockout mice demonstrated a blunted transcriptional response to acute exercise, characterized by the lack of upregulation of well-established exercise responsive transcription factors including *Nr4a3* and *Ppargc1a*. Six weeks of exercise training in muscle-specific BMAL1 knockout mice induced significantly greater and divergent transcriptomic and metabolomic changes in muscle. Surprisingly, liver, lung, inguinal white adipose and heart showed divergent exercise training transcriptomes with less than 5% of ‘exercise-training’ responsive genes shared for each tissue between genotypes.

**Conclusion:** Our investigation has uncovered the critical role that BMAL1 plays in skeletal muscle as a key regulator of gene expression programs for both acute exercise and training adaptations. In addition, our work has uncovered the significant impact that altered exercise response in muscle plays in the peripheral tissue adaptation to exercise training. We also note that the transcriptome adaptations to steady state training suggest that without BMAL1, skeletal muscle does not achieve the expected homeostatic program. Our work also demonstrates that if the muscle adaptations diverge to a more maladaptive state this is linked to increased inflammation across many tissues. Understanding the molecular targets and pathways contributing to health vs. maladaptive exercise adaptations will be critical for the next stage of therapeutic design for exercise mimetics.

## 1. Introduction

Circadian transcription factors and their autoregulatory transcription-translation feedback loops (TTFL) regulate skeletal muscle homeostasis through both transcriptional and epigenetic regulation [1–3]. The daily gene expression program regulated by the clock mechanism, called clock output, contributes to temporal patterning of a wide range of biological processes including skeletal muscle metabolism and proteostasis [4], as we and others have previously published [5–8].

A clear interaction between exercise and the circadian TTFL has been established in recent years. Genes encoding the negative limb of the TTFL have been shown to be responsive to acute exercise/muscle contraction (e.g., *Pers*, *Nfil3*) [9–16] and exercising at different times of the day is known to lead to unique metabolomic and transcriptional outcomes [17–20]. Exercise training also directly impacts the circadian transcriptome, by increasing the number of clock output genes [21,22]. Various exercise performance measures such as strength and endurance capacity also present time-of-day differences, highlighting a likely role of the muscle clock in overall muscle function [21–27]. While it is clear that exercise and the skeletal muscle circadian TTFL are linked [4], the question of whether the muscle clock has any impact on either the acute response or training induced adaptations in skeletal muscle is yet to be elucidated.

Here we asked whether the core clock factor, *Bmal1* is necessary in skeletal muscle for either the acute exercise response and/or adaptations to exercise training. Additionally, we probed the systemic impact of the loss of muscle *Bmal1* on the adaptations in other peripheral tissues. Our results determined that the core clock factor *Bmal1*, is necessary for the acute molecular response to exercise with notable loss of up-regulation of well-known exercise response genes including *Ppargc1a* and *Nr4a3* [28,29]. We also determined that muscle *Bmal1* is required for the expected transcriptional and metabolomic response to treadmill training in muscle. We found less than 5% of the exercise training induced changes in gene expression were identified in the absence of muscle *Bmal1*. Lastly, we found that loss of muscle *Bmal1* also contributes to a significant divergence in the exercise training transcriptional changes in all peripheral tissues studied including the heart, lung, liver, and white adipose tissue. Importantly these divergent transcriptional outcomes with acute exercise and following training were apparent even though the exercise workloads, speed, grade, and duration, were equivalent between genotypes. Our results identify the core circadian clock factor, *Bmal1*, in skeletal muscle as an integral transcriptional modulator for acute exercise and training adaptations. In addition, these results highlight the importance of the muscle for peripheral tissue transcriptome changes in response to exercise training.

## 2. Materials & Methods

### 2.1. Ethical Approval

All animal and experimental procedures in this study were conducted in accordance with the guidelines of the University of Florida for the care and use of laboratory animals (IACUC #202100000018). The use of animals for exercise protocols was in accordance with guidelines established by the US Public Health Service Policy on Humane Care and Use of Laboratory Animals.

### 2.2. Skeletal Muscle Specific *Bmal1* Knockout Mouse Model

Male inducible skeletal muscle-specific *Bmal1* knockout mice were generated as previously described [5,6]. Activation of Cre recombination was performed by intraperitoneal injections of tamoxifen (2 mg/day) dissolved in sunflower seed oil for five consecutive days when the HSA-Cre+/−; Bmal1 fl/fl mice reached ∼4 months of age. These mice are referred to as iMSBmal1KO mice henceforth. Controls were Cre^+/−^:Bmal1^fl/fl^ mice treated with a vehicle (15 % ethanol in sunflower seed oil) and are referred to as Vehicle treated mice.

To assess recombination specificity within the gastrocnemius DNA was extracted from ∼1-5mg of muscle post-mortem, lysed in tail lysis buffer (10mM Tris pH 8, 100mM NaCl, 10mM EDTA, 0.5% SDS and 20mg/ml of Proteinase K) and DNA purified using the DNeasy Blood and Tissue Kit (Qiagen, Venlo, Netherlands). PCR was performed with extracted DNA and primers for the recombined and non-recombined alleles as previously described [5]. For the training study, lack of recombination was confirmed in the inguinal white adipose tissue, liver, heart, and lung. Recombination specificity for tissues in this cohort is provided in Supplementary Figure 1.

A total of 40 mice were used in this experiment, (*n*=20 vehicle and *n*=20 iMSBmal1KO mice). Mice were randomly assigned to a control or an acute exercise group (Vehicle Control *n*=10, Vehicle Exercise *n*=10, iMSBmal1KO Control *n*=10, iMSBmal1KO Exercise *n*=10). Mice assigned to the acute exercise bout experiment were group housed (2-5 per cage) at 23 ± 1.5°C, 58 ± 5.8% relative humidity within temperature and humidity controlled light boxes. Light schedules were maintained on a 12:12 h light: dark schedule with *ad libitum* access to standard rodent chow (Envigo Teklad 2918, Indianapolis, IN, USA) and water. The macronutrient content of this chow is 24% protein, 18% fat and 58% carbohydrate, presented as a percent of calorie intake. Experiments were performed in two cohorts that differed significantly by age (30 weeks vs. 77 weeks, *p*<0.0001) (Supplementary Figure 2A), but were randomly and evenly distributed across the vehicle and iMSBmal1KO groups. Following randomization, no differences in age were observed between ages of vehicle and iMSBmal1KO mice, (*p*=0.86), (Supplementary Figure 2B).

An additional 21 mice were used for exercise training experiments. 21 male mice were randomly assigned to tamoxifen or vehicle treatment, with or without exercise training at ∼11 months of age (7-months post tamoxifen treatment). For clarity, these groups will be referred to in future text as Vehicle Control (*n*=5), Vehicle Exercise Trained (*n*=5), iMSBmal1KO Control (*n*=6), and iMSBmal1KO Exercise Trained (*n*=5). Mice were single housed in conditions of 23 ± 1.5°C, 58 ± 10% relative humidity within temperature and humidity controlled light boxes. Light schedules were maintained on a 12:12 h light: dark schedule with *ad libitum* access to standard rodent chow (Envigo Teklad 2918, Indianapolis, IN, USA) and water. To maintain standardised nomenclature for this study, we will reference time through use of zeitgeber time (ZT), where ZT0 refers to time of lights on and ZT12 refers to time of lights off. The active period of the day for mice is from ZT12-ZT24 and their rest period is ZT0-ZT12. All physiological testing, treadmill familiarization and exercise/training sessions were performed in the active phase so transportation of mice from housing to the treadmill room was in a light tight cart and the treadmill room was dark with red lights.

### 2.3. Treadmill Familiarization, Graded Exercise Test, Grip Strength & Rotarod

Mice from all groups/experiments were first subjected to treadmill familiarization under red light on a Panlab treadmill (Harvard Apparatus, Holliston, MA) at ZT13. Familiarization consisted of 3 sessions performed on 3 consecutive days. The first familiarization session started at 10 cm/s for 5 min at 0° incline. The incline was then adjusted to 5° and speed was increased by 2 cm/s every 2 min up to 20 cm/s (15 minutes total). The second day consisted of an increase in speed every 3 min by 3 cm/s from 10 cm/s to 24 cm/s at an incline of 10°, (18 minutes total). The final familiarization session was the same as the second but performed at a 15° incline, (18 minutes total).

Following 1 day of rest, mice were subject to a graded treadmill exercise test to serve as a baseline performance measure, conducted at ZT13 under red light. For the graded exercise test, animals began at a speed of 10 cm/s at 10° incline for a 5 min warm-up. From which the incline was increased to 15° and the treadmill speed increased by 3 cm/s every 2 min until exhaustion. The treadmill was operated with the electrical shock grid turned off and mice gently encouraged by a compressed air canister when needed. Mice were deemed to be exhausted when they remained in contact with the back of the treadmill >15 s and could not be encouraged to move away by the compressed air.

Mice in groups assigned to an acute exercise bout were given ∼1-week of rest before their acute exercise bout. This consisted of a 30-minute treadmill bout at ZT13 in which the mice would complete 70% of the work done within the graded exercise test (group mean), calculated according to previous work [30], work done = (M⋅D⋅ sin q). M is the mass (g) of the animal, D is the distance (m) ran in the graded exercise test and sin q is the slope (15°) of the treadmill. Work done is measured in arbitrary units (AU).

Mice in groups assigned to exercise training were given 2-weeks of rest post familiarization before their 6-weeks of exercise training commenced. Exercise training was always completed at ZT13 under red light, 5 times per week for a total of 30 individual exercise training bouts. Each bout consisted of 1 hour of treadmill running in which mice would complete 100% of the work done in their prior graded exercise test (group mean). Additional graded exercise tests were conducted at the end of 2 and 4 weeks of training to account for increases in performance and adjustment of exercise speed during training sessions. As a result, treadmill speed was set to 18cm/s for weeks 1-2, 25.5cm/s for weeks 3-4 and 29cm/s for weeks 5-6 for both groups. At the end of 6-weeks training a final maximal capacity test was performed to serve as a post training comparison.

Forelimb grip strength (N/gBW) measures were performed between ZT13-14 on mice from the acute exercise cohort using a Harvard Apparatus grip strength meter (average of 5 trials). Rota Rod (Panlab) performance was also measured in 3 trials and averaged, data is presented in Supplemental Figure 1.

### 2.4. Body Composition Analysis (Echo-MRI)

Body composition was assessed by nuclear magnetic resonance (Echo MRI-100 Body Composition Analyzer; Echo Medical Systems, Houston, TX) prior to treadmill familiarization at ZT13. For the exercise training cohort, measurements were taken in the week prior to starting exercise training, after 3 weeks and after 6 weeks of exercise training. Lean and fat mass (grams) are presented as percentage changes over time as well as a percentage of total body weight over time.

### 2.5. Food Intake & Cage Activity Measurements

As described above, mice were individually housed to track food intake and cage activity. Food and body mass were weighed on a weekly basis to assess changes in food intake relative to body weight. The daily activity of the mice in their home cages were continuously recorded throughout the entire experiment using wireless infrared activity monitoring (Actimetrics, Wilmette, IL, USA) and analysed using ClockLab software 7. Average counts per week were calculated for the 4-weeks prior to training, weeks 1-2, 3-4 and 5-6. The hour during treadmill running +/− 30 minutes was excluded from each individual’s analysis.

### 2.6. Tissue Collection

All animals were humanely sacrificed following the experiments using isoflurane anaesthesia (2-3%) in oxygen followed by cervical dislocation performed under red light. For the acute exercise bout experiments, tissue was collected 1-hour post the end of their acute exercise bout (ZT14.5). Control, unexercised mice were collected alongside the acute exercise groups as time-of-day matched controls. Gastrocnemius muscles from the hindlimbs were dissected, cleaned of excess fat and connective tissue and flash frozen in liquid nitrogen before storage at −80°C. For the training study, animals were humanely sacrificed as above, 47 hours after the exercise trained mice final maximal capacity test at ZT13. Hindlimb muscles, iWAT, liver, heart and lungs were dissected and cleaned of excess fat and connective tissue. The heart and lungs were gently perfused with phosphate buffered saline. Muscles from the left hindlimb were weighed and flash frozen in liquid nitrogen and muscles from the right hindlimb embedded in optimal cutting temperature compound (OCT) before snap freezing in isopentane pre-cooled by liquid nitrogen. Muscles were stored at −80°C until biochemical analysis or cryo-sectioning.

### 2.7. Blood/plasma collection & Analysis

For the training study, blood samples ∼50ul were collected from the tail veins of all mice at ZT13 prior to the maximal exercise test, 48 hours prior to tissue collection. Blood glucose levels were measured at this point using the OneTouch® UltraMini system (OneTouch Solutions, UK). A whole blood aliquot was kept on ice for assessment of glycosylated hemoglobin content (Hba1c%) (#80310, Crystal Chem High Performance Assays, IL, USA) as per manufacturer’s instructions. Endpoint colorimetric measurements were made using the Molecular Devices Spectramax i3X system (Molecular Devices LLC, CA, USA). The remaining blood was collected in EDTA coated plasma tubes before centrifugation at 4°C for 10 mins at 2000RCF. The plasma was frozen and stored at −20°C. Insulin concentration was measured using a commercially available kit (#90080, Crystal Chem High Performance Assays, IL, USA) as per manufacturer’s instructions.

### 2.8. Gastrocnemius Muscle & Liver Glycogen/Triglyceride Content Analysis

Small (∼15mg) snap-frozen pieces from the gastrocnemius muscle and liver (n=5 per group) were used to detect glycogen/triglyceride content. Care was taken to ensure the same location of each tissue was sampled across replicates. For glycogen, tissue samples were weighed and homogenized in 100μl of glycogen hydrolysis buffer in a bullet blender (BBY24M, Next Advance, NY, USA), at speed 10 for two, 2-min sets, at 4°C using 0.5mm stainless steel beads. Homogenates were boiled at 95°C for 5 mins to inactivate enzymes, then centrifuged at 12,000 rpm for 5 mins at 4°C to remove insoluble material. Muscle glycogen concentrations were then quantified from the supernatant using a commercially available kit (Sigma-Aldrich-MAK016) according to manufacturer’s instructions. Glycogen concentrations were normalized to input tissue weight. For triglyceride analysis, tissue was homogenized as above in chloroform: methanol (2:1). Samples were incubated for 30 mins at 20°C with shaking at 1,400rpm. Samples were then centrifuged at 13,000rpm for 30 mins at 20°C. The liquid phase was mixed with 0.9% NaCl and centrifuged at 2,000rpm for 5 mins. Chloroform: Triton X-100 (1:1) was added to the organic phase and solvents evaporated. Triglycerides were then measured using a commercially available kit (Abcam, (ab65336), according to manufacturer’s instructions. Triglyceride concentrations were normalized to input tissue weight.

### 2.9. Fibre Morphology Analysis

Histological samples were transversely sectioned (10µm thick) from the mid-belly of the gastrocnemius, soleus and plantaris muscles and labelled with a cocktail of antibodies against isotype-specific anti-mouse primary antibodies for types IIA (SC-71) and IIB (BF-F3) (1:100) from Developmental Studies Hybridoma Bank (DSHB) (Iowa City, Iowa, USA), and an antibody against dystrophin (1:100, ab15277, Abcam, St. Louis, MO, USA) overnight as previously described [31,32]. Following overnight incubation, sections were washed for 3×5 mins with phosphate buffered saline (PBS) before AlexaFluor secondary antibodies against Rabbit IgG H&L, Mouse IgG1, Mouse IgG2B (Invitrogen/Thermofisher, USA) were added for 2 hours before 3×5 min washes with PBS. Sections were post-fixed in methanol for 5 mins before 3×5 min washes with PBS. Sections were carefully dried to remove excess PBS before mounting in Vectashield Antifade Mounting Medium (Vector Laboratories, UK). For nuclei counting, sections were just labelled with dystrophin and counterstained for nuclei with DAPI. Additional labelling was performed as above using an antibody against embryonic myosin (BF-G6, DSHB). Embryonic myosin positive fibers were counted manually using Image Jv1.53, excluding intrafusal muscle fibers of muscle spindles, and expressed as a percentage of total fibers identified. Manual counting of central nuclei was also performed in this manner. Whole labelled cross sections were imaged at 20x using a Leica TCS SPE microscope equipped with a DFC7000 camera (Leica, Germany). LAS Navigator software was used to generate a merged image of the whole muscle cross-section. Fiber type distribution, overall muscle fibre cross-sectional area (CSA) and total nuclei/myonuclei per fiber was characterised using MyoVision 2.2 automated software [31]. For succinate dehydrogenase (SDH) staining, cryosections were incubated at 37°C for 1 hour with a sodium succinate buffer (50mM sodium succinate, 50mM phosphate buffer and 0.5mg/ml nitroblue tetrazolium) before washing in distilled water and imaged as above. Colorimetric images were converted to greyscale 8-bit image and SDH labelling intensity quantified using the polygon selection tool to select within the muscle cross-section and raw intensity density measurement tool as standard in Image Jv1.53.

### 2.10. RNA Isolation, Library Preparation & RNA-sequencing

For the acute exercise bout experiment, gastrocnemius muscles were powdered under liquid nitrogen and ∼20mg weighed and placed in 500 μl of Trizol (Invitrogen 15596018). Tissue samples were homogenized in a bullet blender using 0.5mm stainless steel beads (BBY24M, Next Advance, NY, USA) for 2 mins at 4°C. RNA was isolated using the RNeasy Mini Kit (Qiagen 74104) according to the manufacturer’s protocol. DNase digest was performed on column using the RNase-Free DNase Set (Qiagen 79254). RNA concentration and purity were assessed by UV spectroscopy at ODs of 260 and 280 nm using a Nanodrop (ThermoFisher One/One UV spectrophotometer, USA). RNA integrity numbers (RIN) were subsequently determined on an Agilent TapeStation 2200 (Agilent, CA, USA), and all samples prepared for sequencing had RIN values of at least 8. Preparation of poly-A tail selected RNA libraries (Illumina mRNA Prep kit (Illumina, Inc. CA, USA)) and whole transcriptome sequencing (Novaseq 6000 (Illumina, Inc. CA, USA)), was conducted by Novogene Co., LTD (Durham, NC) in a 2×150bp format for the acute exercise bout study. A total of 20 samples were sequenced with a read depth of around ∼25 million paired end reads per sample, *n*=5 per group (∼30 weeks of age).

For the exercise training experiment, RNA was extracted and sequenced in the same format as above but was performed by the University of Florida ICBR Gene Expression and Genotyping Core Facility, RRID:SCR_019145 and the University of Florida ICBR NextGen DNA Sequencing Core Facility, RRID:SCR_019152 for gastrocnemius muscle, liver and heart (*n*=5 per group). iWAT and lung, (*n*=3 per group) were completed as above by Novogene Co., LTD (Durham, NC).

### 2.11. RNA-seq Bioinformatic Analysis

FastQ files were imported to Partek® Flow® Genomic Analysis Software (Partek Inc. Missouri, USA) for pipeline processing. Pre-alignment QA/QC was performed on all reads prior to read trimming below a Phred quality score of 25 and adapter trimming. STAR alignment 4.4.1d was used to align reads to the Mus Musculus, mm10 genome assembly. Aligned reads were then quantified to the Ensembl transcriptome annotation model associated with Mus Musculus mm10, release 99_v2. Gene expression was normalized using DESeq2 median of ratios within tissues. Differentially expressed genes (DEGs) were identified through the Partek® Flow® DESeq2 binomial generalized linear model with Wald testing for significance between conditions. In Supplementary Figure 3, we overlapped the queried vehicle exercise responsive DEGs with a previously published BMAL1:CLOCK chromatin immunoprecipitation with sequencing dataset from our lab to identify directly bound genes within murine gastrocnemius muscle (any genomic region, FDR <0.001) [33].

### 2.12. Assay For Transposase-Accessible Chromatin Using Sequencing (ATAC-seq)

A whole gastrocnemius muscle was diced and then homogenized using a Polytron homogenizer in homogenization buffer (Homogenization buffer (10mM HEPES, pH 7.5, 10mM MgCl2, 60mM KCl, 300mM sucrose, 0.1mM EDTA, 0.1% Triton, 1mM DTT, complete with mini protease inhibitor mix). Homogenates were spun at 300rpm for 3 minutes and supernatant containing nuclei moved to a new tube. This was repeated 3 times. The supernatant was then filtered through a 40µm cell strainer before centrifugation at 1000g for 10 minutes. The nuclei pellet was then resuspended in 1.2ml of homogenization buffer, gently triturated and 200µl loaded on top of 2.15M sucrose cushions (10mM HEPES, pH 7.5, 10mM MgCl2, 60mM KCl, 2.15M sucrose, 0.1mM EDTA, 0.1% Triton, 1mM DTT, complete with mini protease inhibitor mix). Nuclei samples were centrifuged at 13,500g at 4°C for 1.5 hours. Samples were aspirated and washed leaving a nuclei pellet, free of cytoplasmic fragments. Nuclei were quality checked using light microscopy (intact nuclei, no blebbing, free of cytoplasmic fragments) and then counted. 50,000 nuclei were used for input into a commercially available ATAC-seq kit which includes library preparation (#53150, Active Motif, Inc., CA, USA). We have previously determined that this isolation technique yields ∼70% myosin heavy chain expressing nuclei through RNA-sequencing, (unpublished data). Libraries were assessed via Bioanalyzer to compare library fragments ranging from 250-1000bp with an oscillation period at ∼150bp. were sequenced by the UF ICBR Nextgen DNA Sequencing facility RRID:SCR_019152 with ∼70 million paired-end reads per sample in a 2×150bp format.

### 2.13. ATAC-seq Bioinformatic Analysis

Fastq files were imported to Partek® Flow® Genomic Analysis Software (Partek Inc. Missouri, USA) for processing. Pre-alignment QA/QC was performed on all reads prior to read trimming below a Phred quality score of 30 and adapter trimming. Alignment was performed using Bowtie2 to the mm10 genome, and peak detection/calling performed using MACS2. Peaks were quantified, normalized using Deseq2 (median of ratios) before differential analysis to identify differentially accessible regions (DARs) between conditions (significant differences in peak heights *P*= 0.01). ATAC peaks mapping to any genomic region were included in the differential analysis. 24.3% of peaks corresponded to the transcription start site, 5.2% aligned to the transcription termination site, 4% to exons, 8.5% to 5’ untranslated regions (UTR), 3.1% to 3’UTR regions, 29.3% to intronic regions and 25.6% to intergenic regions.

### 2.14. Pathway & Motif Analysis

Biological interpretation of filtered/selected gene lists was performed using DAVID [34] to identify associated Kyoto Encyclopedia of Genes and Genomes (KEGG), Reactome and Biological Processes pathways. HOMER (v4.11) motif analysis was performed on DEGs, to identify enrichment of known motifs (6-12bp long) in the gene body and up to 2kb upstream of the transcription start site. Upset plots were created using Intervene, a publicly available software [35].

### 2.15. LC-MS Metabolomics

Gastrocnemius muscle samples (*n*=5 per group) underwent cellular extraction procedures (Southeast Center for Integrated Metabolomics, SECIM). Global metabolomics profiling was performed on a Thermo Q-Exactive Orbitrap mass spectrometer with Dionex UHPLC and autosampler. All samples were normalized by total protein content prior to extraction. Samples were analyzed in positive and negative heated electrospray ionization with a mass resolution of 35,000 at m/z 200 as separate injections. Separation was achieved on an ACE 18-pfp 100 x 2.1 mm, 2 µm column with mobile phase A as 0.1% formic acid in water and mobile phase B as acetonitrile. This is a polar embedded stationary phase that provides comprehensive coverage but does have some limitations in coverage of very polar species. The flow rate was 350 µL/min with a column temperature of 25°C. 4µL was injected for negative ions and 2 µL for positive ions. MZmine (freeware) was used to identify features, deisotope, and align features. All adducts and complexes were identified and removed from the data set. The mass and retention time data was searched against SECIMs internal metabolite library, and known metabolites were mapped to KEGG IDs. Blank feature filtering was performed using inner-quartile range filtering as implemented in the R package MetaboAnalystR (https://github.com/xia-lab/MetaboAnalystR). Missing data were imputed by k-nearest neighbour imputation and peak intensities were normalized sample-wise using sum normalization followed by log^10^ transformation and pareto scaling as implemented R package MetaboAnalystR. Univariate ANOVA was performed using the R package stats (R Core Team 2023) to determine whether there was a significant effect of variable Class on metabolite peak intensity, and post-hoc pairwise comparisons were performed using least-squares means method as implemented in the R package emmeans (R.V. Lenth 2023). Adjusted p-values are adjusted for multiple pairwise comparison testing, not testing of multiple metabolites. A significance threshold was set at an FDR of 5%. Metaboanalyst 5.0 [36] was used to perform enrichment analysis on annotated features against the KEGG metabolite set library.

### 2.16. Statistical Analysis

Unless otherwise stated, physiology data are presented as mean ± standard deviation (SD) and statistical analyses conducted in GraphPad Prism 9.4.1. For multiple comparisons, data were analysed using one, or two factor analysis of variance (ANOVA) followed by Tukey’s multiple comparisons test. Repeated measures ANOVA were performed on measures where the same subjects were measured over time. To assess differences between exercise training groups only, two-tailed independent student t-tests were performed to evaluate statistical differences. All statistically significant thresholds were considered at the level of *P* < 0.05. Corresponding symbols to highlight statistical significance are as follows; \**P* ≤ 0.05, \*\**P* ≤ 0.01, \*\*\**P* ≤ 0.001, \*\*\*\**P* ≤ 0.0001.

### 2.17. Data Availability

RNA-seq data is available in Supplementary File 1 and raw data can be found at the GEO accessions GSE255280 (acute exercise), GSE240653 (gastrocnemius training), GSE253735 (peripheral tissue training). Chromatin accessibility data (ATAC-seq is available in Supplementary File 1 and raw data can be found at GSE255376. LCMS metabolomics data is available in Supplementary File 2.

## 3. Results

### 3.1 Loss of the skeletal muscle clock significantly alters the acute exercise transcriptional response

Previous work has identified that acute exercise performed at different time of the day leads to divergent transcriptional responses [18] suggesting that the acute response to exercise is modulated by the muscle circadian clock. To directly test this, we exposed vehicle treated and iMSBmal1KO mice to a workload-matched 30-minute treadmill exercise bout and performed transcriptional analysis on the gastrocnemius muscles at 1-hour post completion of the bout, (Figure 1A). Physiological phenotyping is available in Supplementary Figure 2. Despite vehicle and iMSBmal1KO mice completing the same amount of treadmill work (Figure 1B), the acute transcriptional response was significantly different in many parameters. First, we noted the iMSBmal1KO muscle exhibited a 48% reduction in the number of exercise responsive genes with 517 genes changed (250 upregulated and 267 downregulated: *q* < 0.05) compared to the 989 genes that changed in the muscle of vehicle mice (600 upregulated and 389 downregulated), (Figure 1C). Only 117 of these genes were common as exercise response genes in both genotypes, (Figure 1D). Within this shared group of exercise responsive genes ‘Biological rhythms’ was the most enriched pathway (Figure 1E), driven by changes in expression of genes in the negative limb of the E-box feedback loop (*Per1*, *Per2*) and RORE element/D-box feedback loop (*Nfil3*) which are still responding to the exercise bout despite the absence of BMAL1, (Figure 1F)[37], previously shown to be downstream of CREB[38]. This observation confirms that the negative limb circadian genes can be directly regulated downstream of the exercise bout in skeletal muscle and do not require *Bmal1* or a functioning TTFL clock mechanism.

**Figure 1:**
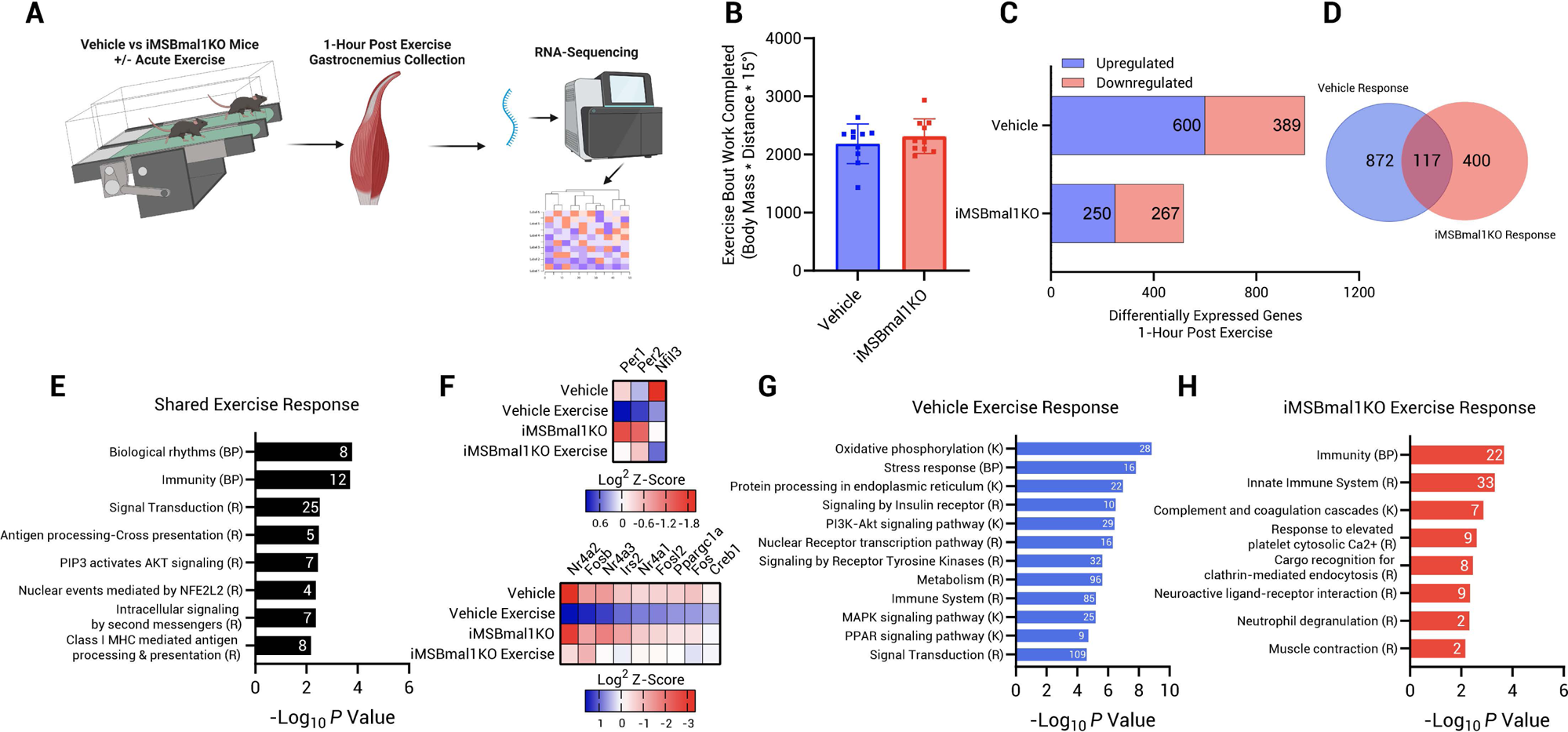
Loss of skeletal muscle *Bmal1* alters the transcriptional response to acute exercise. (A) Experiment 1. Vehicle and iMSBmal1KO mice completed a 30-minute, 15° inclined treadmill bout at 70% of their maximum work done and gastrocnemius muscles were collected 1-hour post. (B) Exercise bout work was matched between genotypes. (C) RNA-sequencing was used to assess the number of DEGs 1-hour post exercise in vehicle and iMSBmal1KO mice. (D) Venn overlap of vehicle and iMSBmal1KO mice exercise responses. (E) Pathway analysis of shared exercise response genes. (F) Z-scores of exercise responsive clock factors and common exercise response genes shown between genotypes. Associated pathway databases are indicated by (R) Reactome, (K) KEGG, and (BP) Biological Processes. Pathway analysis of vehicle exercise responsive genes (G) and iMSBmal1KO exercise responsive genes (H).

Analysis of the exercise DEGs unique to vehicle muscle identified the robust up-regulation of well-known exercise transcription factors and co-activators, including *Nr4a3*, *Fosb* and *Ppargc1a (PGC1ɑ)*. In contrast, this cluster of well-defined exercise responsive factors did not change following exercise in iMSBmal1KO muscle. This was unexpected as these genes have been shown to be very responsive to an acute exercise bouts in preclinical rodent models, in muscle of healthy human subjects and in muscle from subjects with T2D [9,10,28,39], (Figure 1F). These results identify that muscle BMAL1 is necessary for the acute up-regulation of *Ppargc1a*, *Nr4a3* and other exercise responsive transcription factors in skeletal muscle. We suggest that the lack of up-regulation of these factors, in the absence of muscle BMAL1, likely contributes to the divergent transcriptional program post-acute exercise.

We next asked if the divergent exercise gene expression patterns reflected similar or different functional pathways, (Figure 1G-H). Consistent with many studies, the vehicle exercise response pathway enrichment contained ‘Oxidative Phosphorylation,’ ‘Stress Response,’ and signaling pathways such as ‘insulin receptor, ‘PI3K-Akt’, ‘Nuclear Receptor’, Receptor Tyrosine Kinase’, ‘MAPK’ and ‘PPAR’. Additionally, there were 85 ‘Immune System’ associated genes also differentially regulated specifically in vehicle treated mice. Pathway analysis of the iMSBmal1KO specific exercise DEGs revealed very different pathways and noticeably lacked the pathways seen in the vehicle muscle response. The primary pathways identified were enriched for ‘Immunity’ (22 genes) and the ‘Innate Immune System’ (33 genes), (Figure 1H). We noted that 109 ‘signal transduction’ associated genes were altered post exercise, (61% upregulated) in the vehicle mice, but were unchanged in the iMSBmal1KO, (Figure 1G). This included the absence of several well-known exercise responsive signaling pathways including ‘insulin receptor, PI3K-Akt, nuclear receptor, receptor tyrosine kinase, MAPK and PPARG.

Mechanistically, we asked if the altered exercise response in the iMSBmal1KO mice could be due to epigenetic changes after loss of BMAL1. We performed an assay for transposase accessible chromatin with sequencing (ATAC-seq) on gastrocnemius muscle from sedentary vehicle and iMSBmal1KO mice to determine chromatin accessibility. We identified a number of changes in chromatin accessibility between vehicle and iMSBmal1KO mice that may contribute to some of the altered exercise responses, including 1837 more accessible DNA regions and 1973 less accessible regions, (Supplementary Figure 3A-H). We note that ‘Signal transduction’ was the most enriched pathway in the less accessible DNA regions, comprising 152 genes. Furthermore, 28 of those genes also had decreased gene expression suggesting that loss of BMAL1 alters chromatin accessibility in muscle with reduced availability of many signal transduction genes which could be directly impacting their response to exercise, or their upstream regulators. A further 73 ‘Signal transduction’ associated genes were not affected at the chromatin level but were downregulated at the gene expression level with loss of BMAL1 indicating altered transcriptional regulation. We next queried muscle ChIP-Seq data and identified that 31.5% of acute exercise responsive DEGs are bound by BMAL1:CLOCK within the promoter and/or gene body including *Nr4a1*, *Nr4a2*, *Nr4a3*, *Fosb*, *Irs2*, *Fosl2* and *Ppargc1a* which we highlight are unresponsive in the iMSBmal1KO, (Supplementary Figure 3F-H). These observations highlight that the core clock gene, *Bmal1* contributes to both the epigenetic and direct transcriptional network that is a necessary for the acute exercise response in skeletal muscle.

### 3.2 Loss of skeletal muscle BMAL1 does not impact exercise training workloads, body composition, or muscle size and fiber type

We next asked whether muscle *Bmal1* contributes to endurance training adaptations (Supplementary Figure 4-6). We followed a progressive treadmill training approach, performing graded exercise tests before training to set the relative training intensity for each group of mice and then we repeated testing at 2 and 4 weeks to prescribe increases in workload, followed by a post-intervention test at 6 weeks, Figure 2A. Important for interpretations of this training study, we found that the total treadmill work done (AU) across the timecourse of training was not different between genotypes and both groups demonstrated increased performances with training (*P* < 0.0001), (Figure 2B).

**Figure 2:**
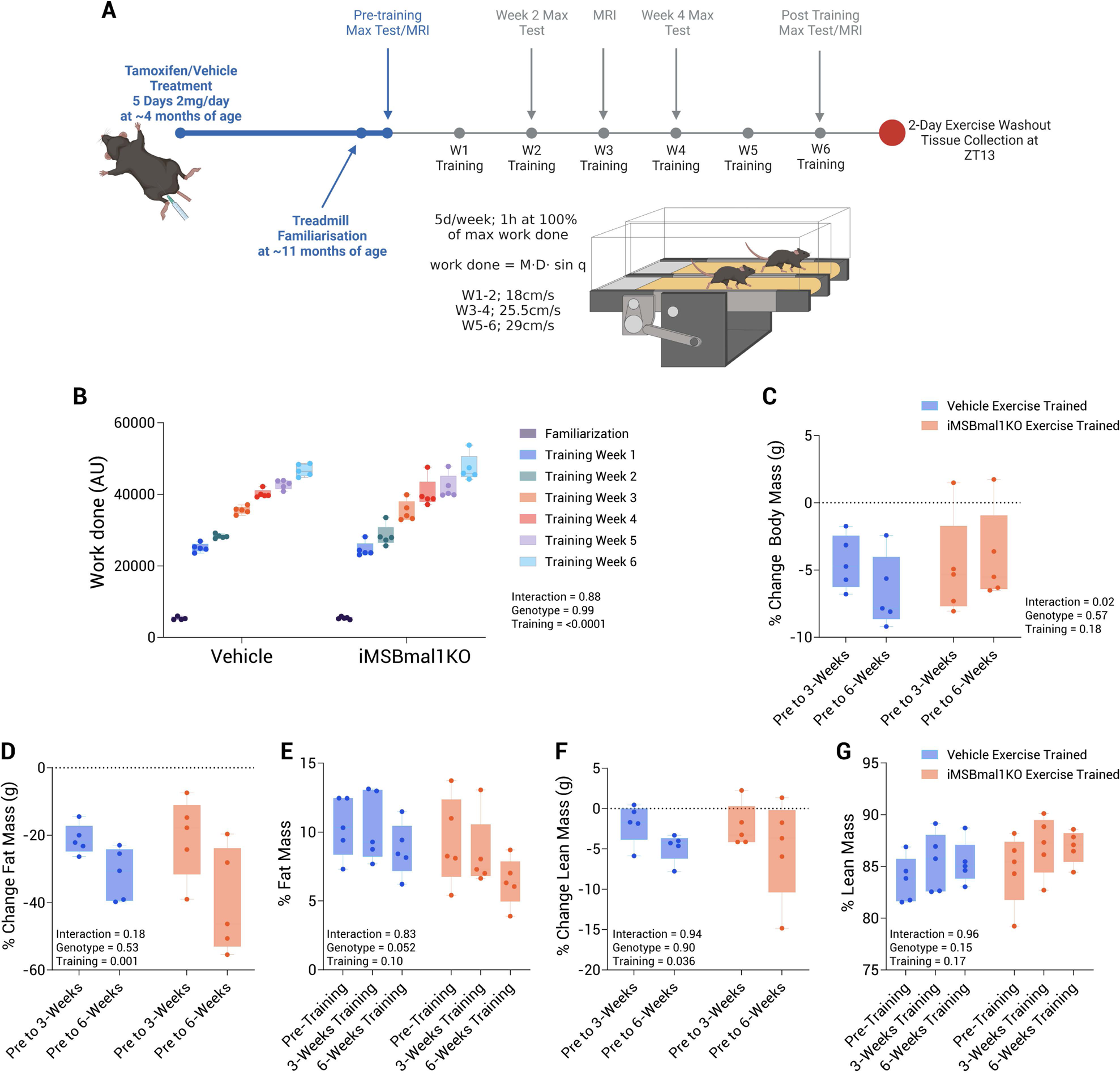
Graded exercise performance and body composition analysis of vehicle and iMSBmal1KO mice. The experimental design is illustrated in (A) which shows the timecourse and parameters of exercise training, physiological testing and tissue collection. At ∼4 months of age, iMSBmal1^fl/fl^ were treated with either a vehicle, or with tamoxifen for 5 days (2mg/day) to induce Cre-recombination and loss of *Bmal1* specifically in skeletal muscle. Both vehicle and iMSBmal1KO mice were randomly assigned to 6 weeks of exercise training performed at ZT13, or control groups. Graded exercise tests were performed prior to, after 2, 4 and 6-weeks of training with the speed adjusted accordingly to make training progressive. MRIs were performed before, after 3 and 6 weeks of training. Mice from all groups were euthanised, and tissue was collected at ZT13, 47 hours after their final graded exercise test. (B) illustrates the weekly work done (AU) during exercise training. (C) Measurements of body mass were taken before, after 3 and 6 weeks of training and the subsequent percentage change calculated for each individual. (D) Total fat mass (g) measured before, after 3 and 6 weeks of training. (E) Percent changes in fat mass (g) were calculated from the beginning of the training period to the middle and end of the training period. (F) Total lean mass (g) measured before, after 3 and 6 weeks of training. (G) Percent changes in lean mass (g) were calculated from the beginning of the training period to the middle and end of the training period.

We observed no differences in body mass between groups, (Supplementary Figure 4C). Following 6-weeks training there was no significant differences between body mass for both of the exercise trained groups vs. their respective controls (Vehicle: 32.3 ± 2.7g vs. 29.8 ± 0.8g, *P* = 0.13 and iMSBmal1KO: 34.1 ± 2.5g vs. 30.5 ± 2.4g, *P* = 0.09), (Figure 2C/Supplementary Figure 4C). Food intake measured per week (g, p/w) per bodyweight (g) did not change over time, or between groups across the experimental timecourse, equaling ∼0.8-1 times bodyweight per week, (Supplementary Figure 4D). We also note that there were no significant differences between the 4 groups in cage activity measures, suggesting that neither loss of muscle *Bmal1* nor exercise training affected behavioral cage activity (Supplementary Figure 4E-G).

Run training has been shown to modify body composition. In this study, MRI body composition measurements identified no significant differences in total fat mass, or percent fat mass as a proportion of total body mass prior to training between genotypes, (Supplementary Figure 4C). We identified a progressive decline in total fat mass in both the vehicle exercise trained (*P* = 0.006) and iMSBmal1KO exercise trained mice, (*P* = 0.03), with no genotype effect (*P* = 0.5), (Figure 2D-G). Similar progressive reductions in lean mass (g) were observed between vehicle exercise trained (*P* = 0.01), equivalent to a 4.8 ± 1.7% reduction in lean mass. A similar percent reduction (4.98 ± 6.1% reduction) was observed in iMSBmal1KO exercise trained mice (Figure 2F), but this was not statistically significant (*P* = 0.07), (Figure 2F). These reductions occurred in the absence of changes in the ratio of lean-to fat mass, (Figure 2D-G).

In general, 6-weeks of run training does not normally induce significant changes in muscle fiber size or fiber type. We performed histological analysis on cross-sections of the gastrocnemius muscle to ask if the muscle of the iMSBmal1KO mice exhibited any unique training induced features. Overall, we found no differences in the main measures of muscle phenotype either in the vehicle mice with training or in the sedentary iMSBmal1KO mice at rest or following training. Supplementary Figure 5. We highlight that there were no significant differences in muscle weight, fiber cross-sectional area, myonuclei per fiber cross-section or proportion of fiber types with either muscle specific *Bmal1* knockout, or exercise training (Supplementary Figure 4A-F). We observed similar increases in SDH staining intensity after exercise training but there was no effect based on genotype, (Supplementary Figure 5D). However, we did notice a small but statistically significant increase in central nuclei (0.92 ± 0.86% fibers vs. 3.9 ± 1.7% fibers, *P* < 0.001) and embryonic positive fibers (0.16 ± 0.13% fibers vs. 1.76 ± 0.84% fibers, *P* < 0.001) after exercise training in the *Bmal1* knockout, which was not present in the vehicle treated group after exercise training, (Supplementary Figure 5G-H). This suggests some small, but reproducible, muscle damage the iMSBmal1KO trained mice. Representative immunohistochemistry images of gastrocnemius muscles are available in Supplementary Figure 5I-J.

### 3.3. Loss of skeletal muscle BMAL1 alters core clock factor gene expression and disrupts the transcriptional regulation of metabolism

To assess the response to training we first had to define, in this cohort of mice, the differential gene expression in the muscles of vehicle control vs. iMSBmal1KO. We identified 1217 DEGs; (*q* < 0.05) between the groups; 535 genes that were upregulated and 682 that were downregulated with muscle specific *Bmal1* knockout respectively, similarly to our reports in Figure 1C. As seen in the volcano plot (Supplementary Figure 6A) and consistent with prior studies [4–6], many of the clock output genes, such as *Dbp*, *Mylk4*, and *Tcap* were significantly down-regulated [40,41]. Motif analysis of the promoters of the 682 down-regulated genes in the iMSBmal1KO determined that there was significant enrichment for E-boxes, and these include several circadian clock (CLOCK, NPAS2) specific E-boxes, as well as MYC specific E-box sequences. Other represented motifs in the down-regulated genes include MEF2 or MADS elements and sites for NFAT which are well recognized pathways in skeletal muscle, (Supplementary Figure 6B). Subsequent KEGG pathway analysis of the DEGs confirmed significant dysregulation of genes associated with ‘circadian rhythms’ (14 genes), as well as a large number of pathways relating to substrate metabolism, (Supplementary Figure 6C). Within this pathway, we identified dysregulation of transcripts essential for glucose and glycogen homeostasis including (*Hk2*, *Rab10*, *Slc2a3*, *Slc2a4*, *Tbc1d1*, *Aldoc*, *Fbp2*, *Gbe1*, *Pgm1*, *Pgm2*, *Phka1*, *Pygm*) as previously reported [4–6].

### 3.4. Identification of BMAL1-dependent and BMAL1-independent transcriptional adaptation to 6-weeks of exercise training in gastrocnemius muscle

While there were few phenotypic differences between the muscles of trained vehicle and iMSBmal1KO mice, we did find that the transcriptional adaptations to training were largely divergent. For this analysis, we performed RNA-seq on gastrocnemius muscle 47 hours after their last exercise bout. Analysis of the transcriptome in muscle from vehicle control and vehicle trained mice identified only 142 DEGs after training which is consistent with prior studies with up-regulation of *Fabp1* and down regulation of *Hadha* [10,42], Figure 3A. Enrichment of DNA binding motifs within the promoter or gene body of the small number of upregulated DEGs included MEF2C, SIX1, Androgen receptor half sites and NFAT, (Figure 3B). Downregulated DEGs were also enriched for DNA motifs relating to EGR1, SMAD4 and NFAT, suggesting the latter contributes to both activation/suppression of exercise DEGs. Pathway analysis revealed the expected exercise adaptations pathways including ‘PPAR signaling’ (11 genes), ‘protein digestion and absorption’ (8 genes), ‘fatty acid metabolism/degradation’ (7 genes), ‘ECM-receptor’ (7 genes), ‘focal adhesion’ (5 genes), ‘calcium signaling’ (7 genes), ‘PI3K-Akt signaling’ (5 genes) and ‘HIF-1 signaling’ (5 genes), Figure 3C. As expected, a number of these DEGs included fatty acid metabolism regulators (*Hadha, Hadhb, Acaa2, Cpt2, Acsl1, Scd2, Acadm, Cd36, Fabp1, Apoa1 and Apoa2*).

**Figure 3:**
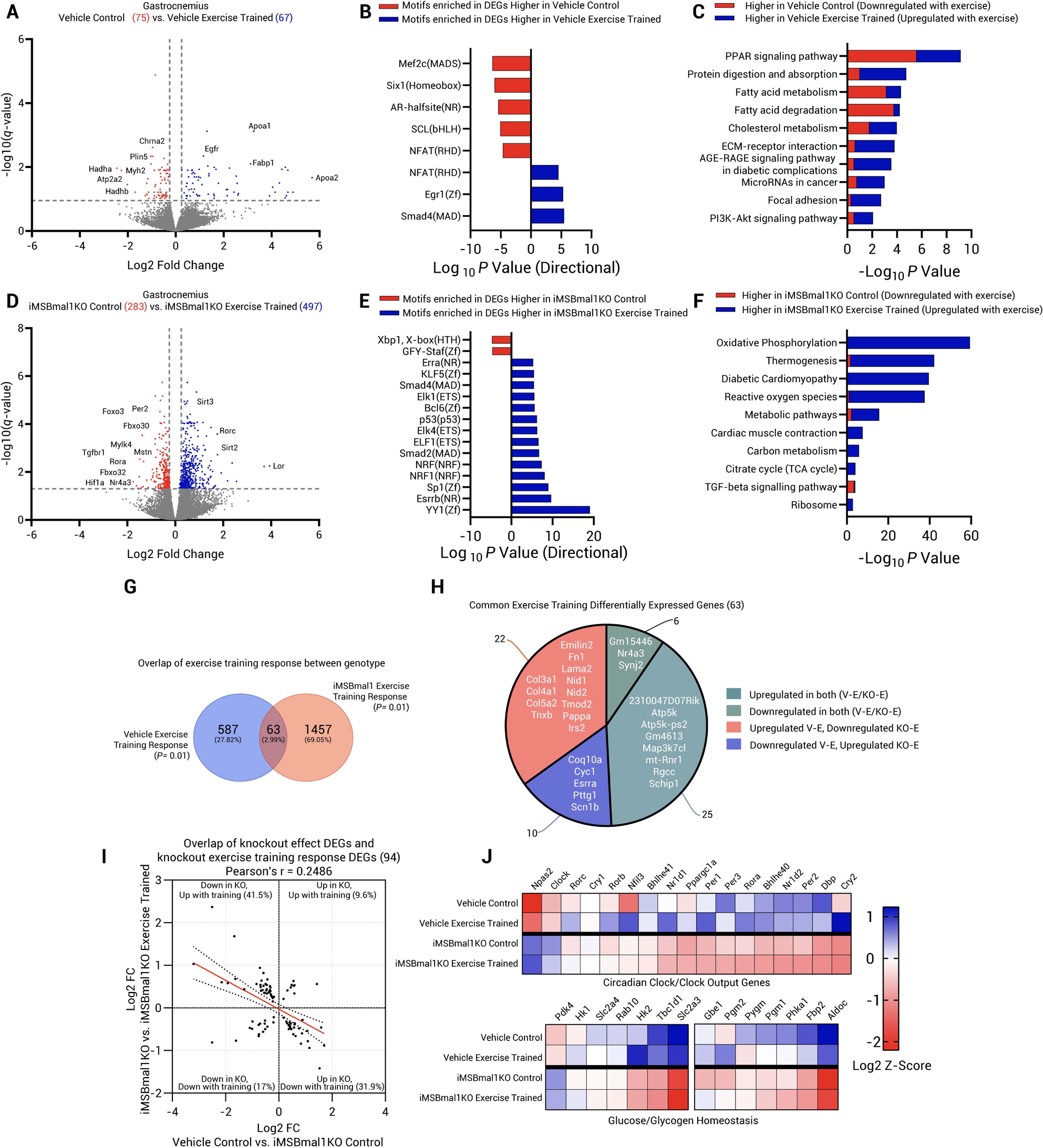
iMSBmal1KO gastrocnemius muscles present a larger and divergent transcriptional adaptation to 6-weeks of exercise training. (A) Volcano plot illustrating DEGs between control and iMSBmal1 WT exercise trained mice. A total of 142 (75 downregulated, 67 upregulated) genes were differentially expressed using an FDR cut off of <0.05. (B) HOMER Motif analysis performed on up and downregulated DEGs identifies a number of transcription factors regulating the transcriptional response to normal exercise adaptation. (C) KEGG Pathway analysis of genes associated with the vehicle treated transcriptional response to exercise training. (D) Volcano plot illustrating DEGs between iMSBmal1KO control and iMSBmal1KO exercise trained mice. A total of 780 (283 downregulated, 497 upregulated) genes were differentially expressed using an FDR cut off of <0.05. (E) HOMER Motif analysis performed on up and downregulated DEGs identifies a number of transcription factors regulating the transcriptional response to exercise adaptation in the absence of skeletal muscle Bmal1. (F) KEGG Pathway analysis of genes associated with the iMSBmal1KO transcriptional remodeling to exercise training. (G) Venn diagram overlapping of DEG responses to vehicle and iMSBmal1KO exercise training at *P* <0.01. Minimal overlap in the transcriptional exercise responses is present between vehicle treated and iMSBmal1KO mice using a lenient significance cut off (63 out of 2107, 2.99% overlap). (H) Pie chart of the 63 common exercise training responsive genes in both genotypes reveals common and oppositely regulated gene reprogramming. (I) Analysis of DEGs affected by loss of BMAL1 and exercise training responsive DEGs in the iMSBma1KO mice, reveals 94 common genes. Pearson’s correlation of the 94 common DEGs (Log^2^ fold changes) reveals that genes affected with Bmal1KO are not rescued with exercise training. (J) Heatmap presenting Log^2^ Z-scores for core clock factors/clock output genes and genes associated with glucose handling and glycogen utilization. Exercise training does not rescue the downregulation of these genes in iMSBmal1KO mice.

In contrast, analysis of the gene expression changes with training in the iMSBmal1KO mice identified a much larger, >5-fold increase in the number of DEGs in the KO muscle following training. We identified 780 DEGs, of which 283 were downregulated and 497 upregulated with exercise training, Figure 3D-F. Similar to our findings following acute exercise, there was little overlap in the exercise DEGs between genotypes. Analysis of the DNA motifs associated with the iMSBmal1 exercise DEGS predicts that exercise adaptation in the iMSBmal1KO muscle relies on vastly different sets of upstream transcription factors, (Figure 3E). Within the genes upregulated by exercise training in the iMSBmal1KO mouse, motif enrichment analysis identified sites for YY1, ESRRA/B, SP1, NRF/1, SMAD2/4 and P53. In the down-regulated genes, we identified XBP1 sites were enriched which was interesting as XBP1 is a transcription factor that functions as a regulator of the unfolded protein response and is known as a key part of the ER stress pathway. For the training study, we collected tissues at 48hrs after the last exercise bout so the observation that many of the DEGs have regulatory elements that indicate a stress pathway was unexpected. We also noted that several of these motifs represent transcription factors normally associated with an acute response to exercise (e.g., ESRRA, NRF1, P53). However, they are not commonly identified as motifs enriched in steady state training induced changes in gene expression [10,43–48], (Figure 3E). The results from the DEG motif analysis suggest a vastly different transcriptional mechanism for muscle adaptation with exercise training that relies on inclusion of known acute response pathways. We suggest that without muscle BMAL1, exercise training results in activation of a chronic stress set of pathways coupled with an inability of the muscle to return to homeostasis following exercise.

While the exercise adaptations were largely divergent, there was a small overlap in the transcriptomes with training between genotypes. Initial analysis identified only 23 common exercise adaptation genes (1.6% of DEGs) using an FDR cut-off of *q* = 0.05. To increase the number of genes for this analysis, we decreased the stringency to *P* = 0.01 and found 63 overlapping genes (2.99%), (Figure 3G). We probed the 63 common genes to ask if they represented a common, non-BMAL1 dependent, training response, Figure 3H. We determined that only 25 (39.7%) of the 63 genes were upregulated in both genotypes, with 6 (9.5%) were downregulated in both. We identify these as BMAL1-independent exercise training genes. Interestingly, 22 (34.9%) of the shared genes were up in vehicle but down in the KO exercise trained muscle and these were associated with the extracellular matrix (*Col3a1, Col4a1, Col5a2, Tnxb, Emilin2, Fn1, Lama2, Nid1* and *Nid2*) and insulin regulation (*Pappa* and *Irs2*). We also identified 10 genes (15.9%) including *Coq10a, Cyc1* and *Esrra* that where downregulated with training in the vehicle but were upregulated in the iMSBmal1KO, (Figure 3H). Thus, while there are shared genes between the two genotypes, over 50% of the 63 shared genes are regulated in opposite directions. This indicates this set of shared genes are exercise responsive, but the directionality is dependent muscle BMAL1.

We also asked whether exercise training can function to restore the dysregulated gene expression in the iMSBmal1KO muscle toward levels seen in vehicle control muscle. For this analysis we compared the DEGs dysregulated by loss of BMAL1 in sedentary mice (1217) to the exercise training DEGs in the iMSBmal1KO muscle (780) and found only 94 DEGs are shared suggesting that the majority of disrupted transcripts in the sedentary iMSBmal1KO muscle are not rescued by exercise training. To visualize this, we plotted Log^2^ fold changes of these genes comparing vehicle control and exercise trained iMSBmal1KO muscle and calculated Pearson’s correlation coefficient. As seen in Figure 3I we identified that 73.4% of the genes disrupted with initial loss of *Bmal1* were directionally closer to vehicle control levels following training. However, the correlation coefficient of determination was weak (R^2^=0.24) indicating that the levels of gene expression were not close to sedentary values. This provides additional evidence that the exercise adaptations work with BMAL1 and are not occurring through separate transcriptional pathways. A heatmap for all 94 genes is presented in Supplementary Figure 7A. Recent results from cardiac *Bmal1* knockout mice have demonstrated that cardiac dysfunction caused by loss of BMAL1 cannot be attenuated with a similar exercise training paradigm [49].

Lastly, we wanted to ask if exercise training impacted core circadian clock genes or genes associated with glucose and glycogen metabolism in muscle. As illustrated in the heatmap in Figure 3J, 6 weeks of training in vehicle mice resulted in changes in many circadian clock related genes such as *Nfil3*, *Per1, Per3*, *Cry2*, and *Dbp*. In contrast, in the iMSBmal1KO mouse there were no changes in any of the factors following training. Since we found that acute exercise can modify expression of clock genes, like *Per1* and *Nfil3*, in the absence of muscle Bmal1 (Figure 1E), the lack of changes in these circadian clock genes with training suggests that it is BMAL1 and the core clock mechanism, that is required for the clock factor changes with training. Analysis of glucose and glycogen metabolism genes found that several of these genes changed expression with training in the muscle, including increases in *Hk2*, *Pgm2* and decreased expression of *Slc2a4* (Glut4). In contrast there were little to no significant changes in gene expression of this cluster of genes in iMSBmal1KO mice post-training. These observations reinforce that BMAL1 and/or a functional circadian clock is required for regulating expression of genes critical for glucose homeostasis in skeletal muscle. Plots with individual data points are provided within Supplementary Figure 8A.

### 3.5. Elevated gene expression of TCA-cycle components, β-oxidation, mitochondrial constituents and mito-ribosomes is a distinct feature of iMSBmal1KO exercise adaptation in gastrocnemius muscle

Analysis of the unique training genes for the iMSBmal1KO muscle identified oxidative metabolic adaptations were a highly enriched category, well beyond any oxidative changes in the vehicle muscle adaptation. We found that a number of these DEGs are shared across pathways relating to pyruvate metabolism, the tricarboxylic acid cycle (TCA cycle), fatty acid β-oxidation and genes relating to mitochondrial complex constituents and their associated mitochondrial ribosomal subunits. A schematic of these pathways is provided in Figure 4A, and the changes in genes within each pathway are presented in Figure 4B-C. It is striking that most genes were upregulated in response to exercise training in the iMSBmal1KO mice, in contrast, muscle from vehicle treated mice showed limited changes for these genes in response to training.

**Figure 4:**
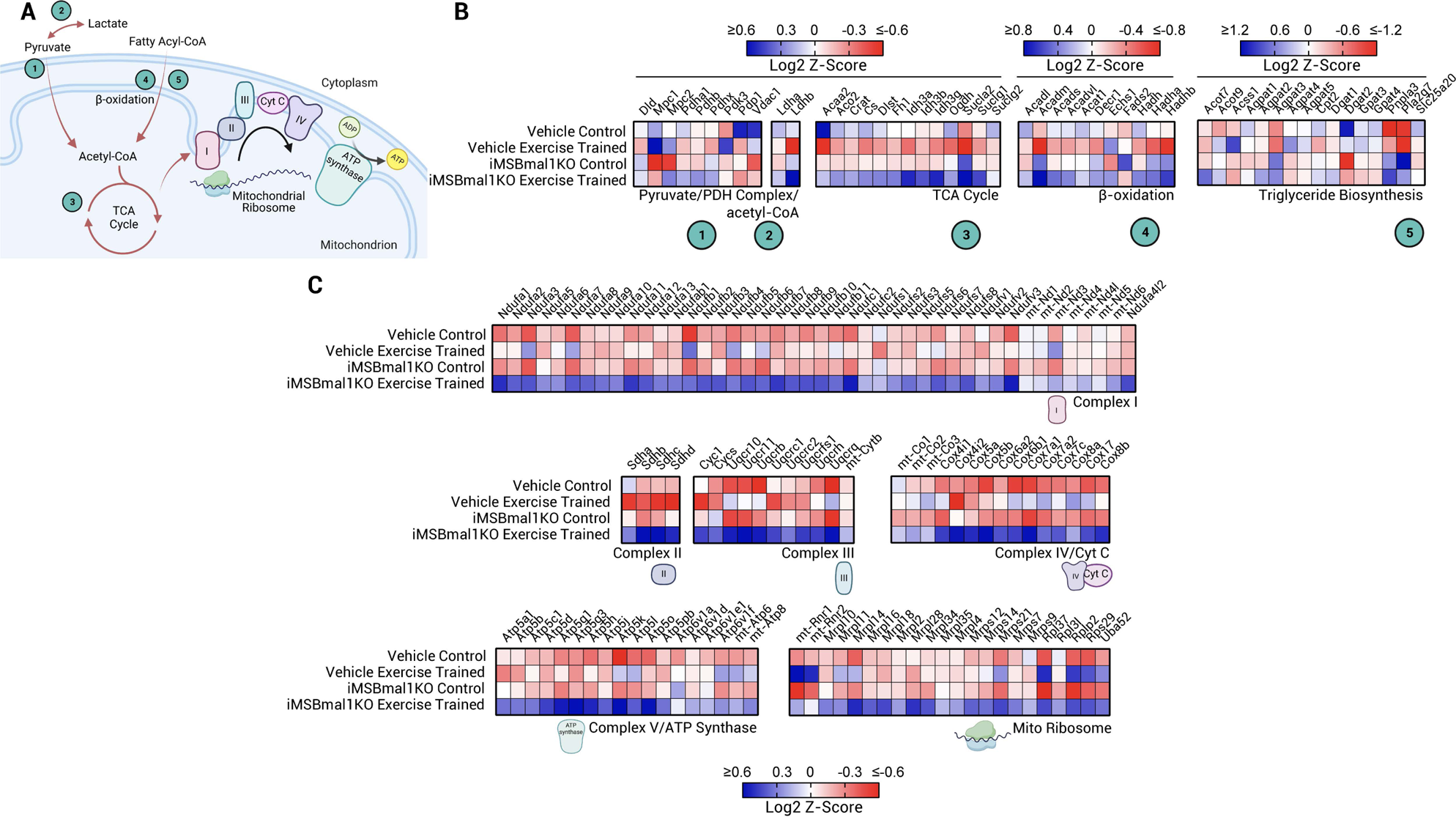
Upregulation of genes associated with pyruvate oxidation, TCA cycle, β-oxidation, mitochondrial complexes and mitoribosome subunits after exercise training in the iMSBmal1KO mouse. (A) Schematic of biological processes involved in providing acetyl-CoA from carbohydrates and lipids for the TCA cycle. Subsequently, the TCA cycle provides NADH and FADH_2_ for the electron transport chain. Components are numbered/labelled for clarity of the associated genes presented in panels B and C. (B) Heatmaps presenting genes associated with pyruvate oxidation, TCA cycle and β-oxidation, (Log^2^ Z-scores). (C) Heatmap presenting genes associated with mitochondrial complexes I-V and the mitochondrial ribosome (Log^2^ Z-scores).

Most strikingly in the iMSBmal1KO training response was the robust up-regulation across almost all genes relating to the TCA cycle, β-oxidation and mitochondrial components encoded by both nuclear and mitochondrially encoded genes, (Figure 4B-C). Interestingly, 12 genes (*Acaa2, Aco2, Crat, Cs, Dlst, Fh1, Idh3a, Idh3b, Idh3g, Ogdh, Suclg1 and Suclg2*) encoding proteins involved in TCA cycle intermediate and anaplerotic reactions were significantly upregulated in the iMSBmal1KO mouse in response to exercise training, whereas they were downregulated with training in the vehicle treated mice. This result suggests that exercise training without muscle BMAL1 pushed increased reliance on alternate substrates to support the energy demand of exercise training. This trend was also observed for a large family of genes associated with Acyl-CoA hydrolysis to fatty acids and coenzyme A and β-oxidation. Genes including *Acadl, Acadm, Acads, Acadv1, Acat1, Acot7* were largely down regulated with exercise training in the muscle of vehicle treated but were uniformly upregulated with training in the iMSBmal1KO muscle. This observation is consistent with a greater requirement for the use of fatty acids for energy metabolism, Figure 4B. We note that there were very few changes in genes involved in triglyceride biosynthesis with exercise training in the iMSBmal1KO indicating that the primary adaptation with exercise was to enhance fat oxidation.

We also observed upregulation of a large network of both nuclear and mitochondrial encoded genes that were largely unique to the iMSBmal1KO training adaptation. These related to mitochondrial complexes I (*n*=44), II (*n*=4), III (*n*=11), IV (*n*=15), V (*n*=18), as well as 22 nuclear and mitochondrial encoded genes related specifically to the large and small mitochondrial ribosomal subunits, (Figure 4C). The majority of these genes were significantly upregulated relative to both vehicle control and vehicle exercise trained mice, as well as iMSBmal1KO control mice. The uniformity of this training response suggests a common transcriptional program for these genes in the absence of BMAL1. Analysis of upstream motifs within these genes identifies sites for *Esrra, Esrrb, Nrf/1* and *p53* and we did identify that expression of *Esrra* was elevated with training in the iMSBmal1KO but downregulated with vehicle exercise trained muscle. As noted earlier, these transcription factors are most seen with the acute exercise response in muscle but not normally associated with steady state training responses, (Figure 3E).

### 3.6. Exacerbated amino acid availability, further depletion of TCA cycle intermediates suggests inefficient energy status with exercise training in the iMSBmal1KO mouse

Transcriptomic data indicated divergent metabolic programs in the muscle of trained vehicle vs. trained iMSBmal1KO mice. We performed untargeted metabolomics to assess potential steady state changes in muscle metabolites following training in the vehicle vs. iMSBmal1KO mouse. Comparing vehicle controls to iMSBmal1KO sedentary mice, we found 46 differentially abundant metabolites, (Figure 5A), a comparison that has previously been reported [25]. In response to training, we observed only 19 differentially abundant known metabolites in vehicle exercise trained mice, which is similar to the small transcriptomic response. In contrast, we identified 116 differentially abundant metabolites in the iMSBmal1KO exercise trained mice. Comparisons between genotypes identified only 8 common metabolites in the exercise training response and they showed enrichment in lysine degradation and glutathione metabolism pathways, (Figure 5B).

**Figure 5:**
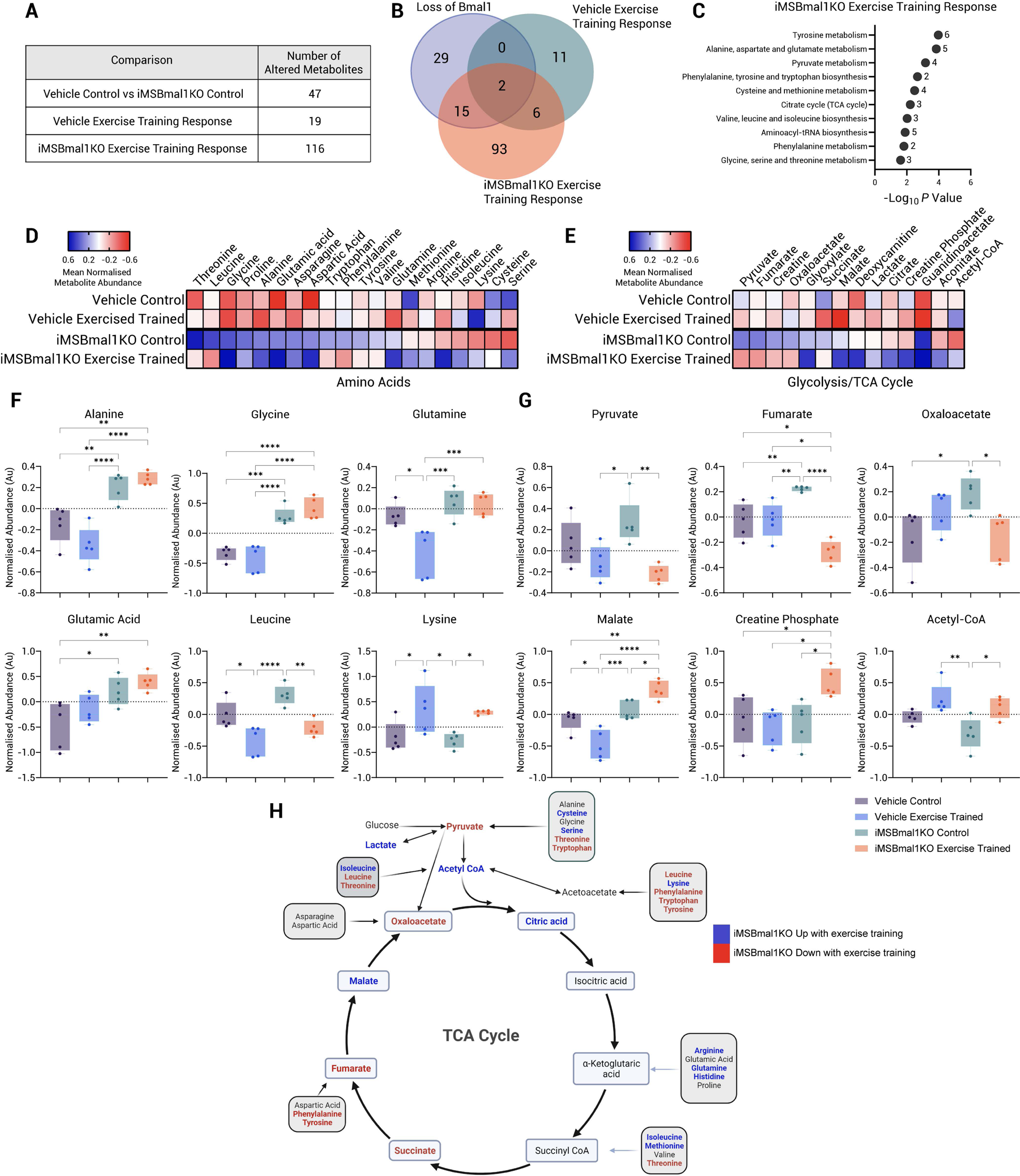
Loss of *Bmal1* in skeletal muscle disrupts the metabolomic rewiring in response to exercise training and causes depletion of TCA cycle intermediates. (A) Table of differentially abundant metabolites (*q* < 0.05) with loss of skeletal muscle BMAL1, the vehicle exercise training response and the iMSBmal1KO exercise training response. (B) Venn diagram analysis of the different comparisons reveals little overlap and divergent metabolomic adaptations to exercise training. (C) Metaboanalyst KEGG metabolite pathway analysis of the iMSBmal1KO exercise training response. Numbers indicate the number of metabolites identified within the pathway. (D) Heatmap illustrating normalized metabolite abundance for amino acids. (E) Heatmap illustrating normalized metabolite abundance for metabolites relating to glycolysis and the TCA cycle. (F) Boxplots of highlighted amino acids changing with loss of BMAL1 and exercise training. (G) Boxplots of highlighted TCA cycle metabolites/phosphocreatine changing with loss of BMAL1 and exercise training. (H) Summary diagram of TCA cycle metabolites affected with exercise training in the iMSBmal1KO (Blue indicates higher abundance, red indicates lower abundance. Significance for one way ANOVA analysis is presented as \**P* ≤ 0.05, \*\**P* ≤ 0.01, \*\*\**P* ≤ 0.001, \*\*\*\**P* ≤ 0.0001.

The differentially abundant metabolites in the iMSBmal1KO exercise trained muscle were enriched amongst amino acid and TCA cycle pathways including ‘Tyrosine metabolism’, ‘Alanine, aspartate and glutamate metabolism’, ‘Pyruvate metabolism’, (Figure 5D-H). Since exercise training does not restore glucose metabolic genes in the iMSBmal1KO muscle we suggest that these metabolite changes reflect the need for TCA cycle substrates via amino acid metabolism. For example, threonine, leucine, tryptophan, phenylalanine and tyrosine are significantly elevated with loss of skeletal muscle BMAL1 but levels were reduced following exercise training. Consistent with this pattern, fumarate, a TCA cycle intermediate metabolized from aromatic amino acids is also significantly upregulated with loss of BMAL1 in sedentary mice and is significantly reduced following training to levels below vehicle controls, (Figure 5E/G). The changes in these metabolites are consistent with the increased reliance on amino acids to support TCA metabolism in the iMSBmal1 muscle with exercise training.

We were surprised that despite our samples being taken 47 hours after their last exercise bout, pyruvate levels were significantly lower by ∼180% in the iMSBmal1KO mice with exercise training, (Figure 5G). This finding falls in line with our transcriptome analysis, that the genes encoding the enzymes required for glucose transport and metabolism are not rescued with exercise training, which may lead to pyruvate availability being a limiting factor. However, we noted above that there is a compensatory increase in genes involved in pyruvate metabolism and TCA cycle components, likely resultant of diminished steady state levels of pyruvate with training. Despite the reduction in pyruvate, acetyl-CoA levels were significantly increased in the iMSBmal1KO mice after exercise training, suggesting that other substrates (likely lipid) contribute more to the increase in acetyl-CoA levels.

Further evidence of disrupted glycolytic metabolism was found when we measured tissue glycogen content in gastrocnemius and liver with exercise training, Supplementary Figure 9D-G. Vehicle treated mice had greater concentrations (*P* = 0.008) and achieved larger increases in muscle glycogen with exercise training (262.8% vs. 66.5%, 1.85 ± 0.13µg/mg vs. 1.43 ± 0.31µg/mg). Similarly in liver, training significantly increased glycogen content in both vehicle treated (16.28 ± 1.81µg/mg, *P* < 0.0001) and iMSBmal1KO mice (13.26 ± 1.64µg/mg, *P* < 0.0001), yet glycogen content was significantly lower in the trained iMSBmal1KO liver, (*P* = 0.036). As the magnitude of improvement in muscle and liver glycogen with training are reduced, it suggests ongoing glycolytic dysregulation within iMSBmal1KO muscle requires higher triglyceride utilization with a systemic impact on the liver glycogen stores.

Glycosylated haemoglobin was lower post exercise training in both groups, *P* < 0.001, with no detected genotype effect, (Supplementary Figure 9A-B). Exercise training significantly reduced plasma insulin levels in both genotypes, however the levels in the exercise trained iMSBmal1KO mice remained significantly higher than vehicle control (∼41%, *P* = 0.0196) and vehicle exercise trained mice (∼135%, *P* <0.0001), Supplementary Figure 9C. While these circulating markers of glucose homeostasis are improved with exercise training in the iMSBmal1KO mouse we note that the effect is partial. We suggest that these partial systemic improvements in blood glucose and insulin are mediated by other peripheral tissue adaptations, such as liver and iWAT, to exercise training.

### 3.7. Loss of skeletal muscle BMAL1 disrupts the transcriptome of peripheral tissues

Exercise is well recognized to induce healthy systemic adaptations and virtually all bodily tissues exhibit significant transcriptome changes following exercise training [50,51]. Since our exercise workloads were the same between genotypes across training, we asked if the peripheral tissue adaptations to exercise would be independent of the divergent muscle adaptations in the iMSBmal1KO mice. For this analysis we chose to investigate highly metabolic tissues and organs that are known to be important for exercise capacity (iWAT, liver, heart and lung). We defined the transcriptomic changes in the vehicle vs. iMSBmal1KO mouse tissues to allow for clear delineation of the exercise training responses. In the control iMSBmal1KO mice we identified a large number of genes (1029) specifically dysregulated in the heart, 511 in iWAT, 159 in liver and 87 DEGs in the lung illustrating unique tissue disruption in gene expression with loss of muscle BMAL1. We did not identify any DEGs shared across all 5 tissues in response to iMSBmal1KO, but we found minor overlap with DEGs specifically dysregulated in muscle and heart (81 genes), muscle and iWAT (58 genes), muscle and liver (21 genes), Figure 6A.

**Figure 6:**
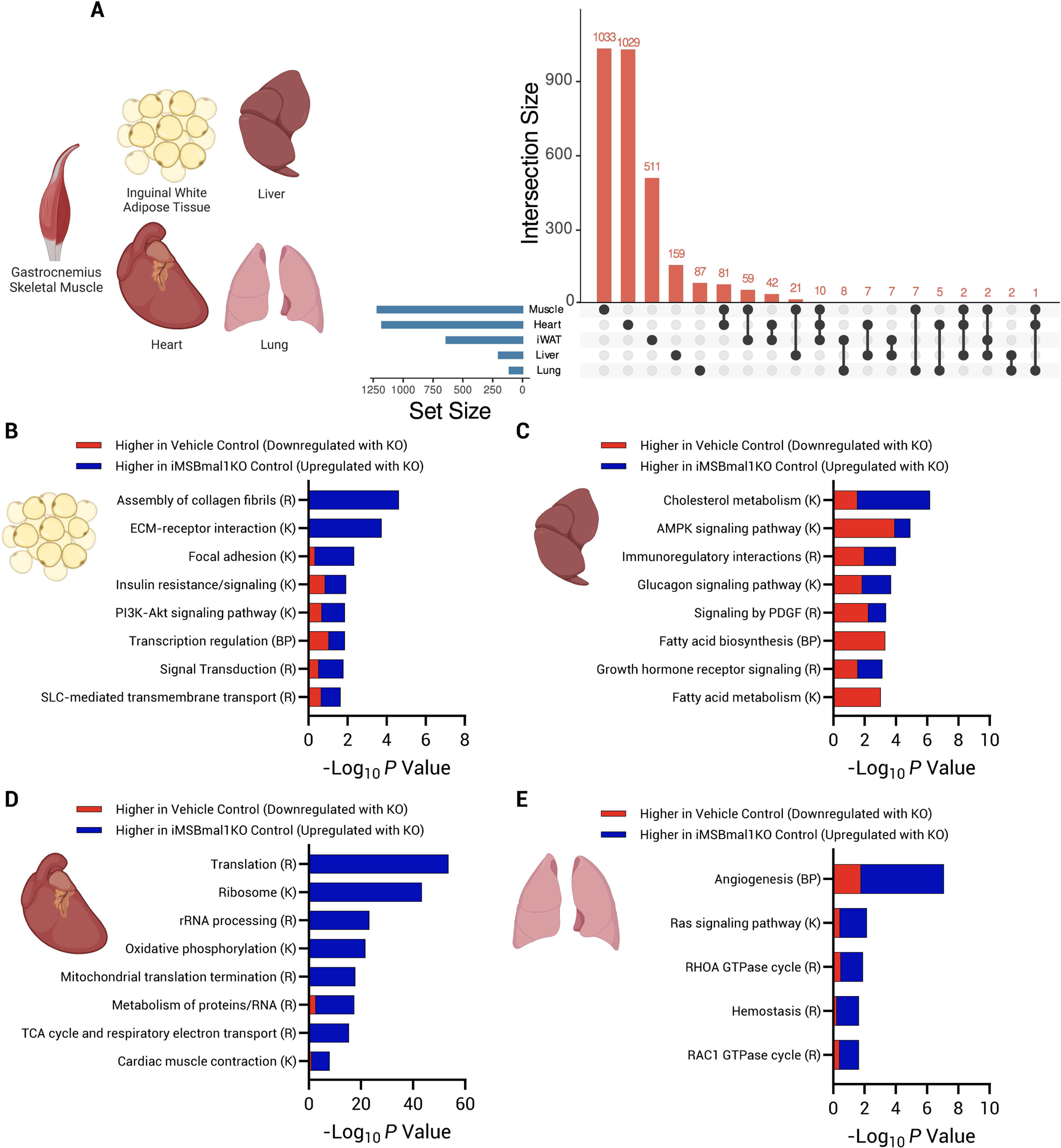
Loss of skeletal muscle BMAL1 indirectly disrupts transcriptional homeostasis of peripheral tissues. (A) Upset plot of the differentially expressed genes affected by loss of skeletal muscle BMAL1 across tissues. Set size indicates the total number of differentially expressed genes regulated by loss of BMAL1 in skeletal muscle in each tissue. Numbers above bars indicate the number of differentially expressed genes. Overlapping differentially expressed genes which are common between tissues are indicated by the connected points below bars. In summary when comparing vehicle control and iMSBmal1KO controls we identified 684 (425 upregulated, 259 downregulated) differentially expressed genes in inguinal white adipose tissue (iWAT), 199 (86 upregulated, 114 downregulated) in liver, 1177 (665 upregulated, 512 downregulated) in the heart and 110 (82 upregulated, 28 downregulated) in lung tissue where identified. Pathway analysis for differentially expressed genes in inguinal white adipose tissue (B), liver tissue (C), heart tissue (D) and lung tissue (E). Red segments of bars indicate downregulation with loss of BMAL1, blue segments of bars indicate upregulation with loss of BMAL1. Associated pathway databases are indicated by (R) Reactome, (K) KEGG and (BP) Biological Processes.

In iWAT, loss of muscle *Bmal1* was associated with 684 DEGs (425 upregulated, 259 downregulated) that were associated with pathways relating to collagen/extra-cellular matrix terms and insulin signaling/resistance related terms which were largely upregulated. *Srebf1*, a key transcription factor responsible for regulation of de novo lipogenesis and glycolysis, as well as *Acacb* which encodes the gene responsible for the carboxylation of acetyl-CoA to malonyl-CoA, a rate-limiting step in fatty acid synthesis and pro-glycogenic genes, *Gys1* and *Ppp1r3d* were significantly downregulated in the iWAT with loss of skeletal muscle BMAL1. This was concomitant with significant upregulation of *Creb3l1*, a master regulator of fibrosis in adipose and liver tissue [52], changes that have previously been associated with insulin resistance and obesity [53–55], (Figure 6B).

The liver transcriptome was also significantly affected by loss of skeletal muscle BMAL1 with 199 DEGs (86 upregulated, 114 downregulated) associated with dysregulation of ‘fatty acid/cholesterol metabolism’, ‘AMPK signaling’ and ‘glucagon signaling’, (Figure 6C). This included significant downregulation of several genes regulating lipid transport/homeostasis and glycolysis including *Elovl6*, *Scd1*, *Apoa4*, Pfkfb1, *Slc2a4* and fatty acid synthesis rate limiting enzyme, *Acaca*. Conversely, *Abcg8*, *Lrp1*, *Lipg* were significantly upregulated along with *Pparg*, known as a master regulator of lipogenesis and glucose uptake [56–59], suggesting a shift of fuel preference to fatty acid metabolism in the liver.

Surprisingly, a large remodeling of the transcriptome was identified in the heart with alteration of 1177 genes (665 upregulated, 512 downregulated). We confirmed that there was no *Bmal1* recombination in the heart tissue of these mice, Supplementary Figure 3. These transcriptional changes included upregulation of genes associated with ribosomal, mito-ribosomal and mitochondrial constituents, proteasomal subunits, but also genes associated with cardiac muscle structure and function. This included upregulation of *Myl2*, *Myl3*, *Tnnc1*, *Tnni3*, *Tpm4*, *Actb* and downregulation of *Dmd*, *Lama2*, *Ryr2*, and *Ttn*, (Figure 6D). More minor changes in gene expression were identified in the lung with 110 DEGs (82 upregulated, 28 downregulated) largely relating to angiogenesis and Ras signaling, (Figure 6E).

### 3.8. Exercise training adaptations are altered in peripheral tissues in mice lacking skeletal muscle BMAL1

We next tested whether exercise training adaptations in peripheral tissues were impacted in the iMSBmal1KO mice. As evidenced by the upset plot, (Figure 7A), we found that all peripheral tissues studied in the iMSBmal1KO mouse exhibited significantly larger numbers of post exercise training DEGs (iWAT 441 vs. 455 DEGs, Liver 336 vs. 571 DEGs, Heart 261 vs. 798 DEGs and Lung 208 vs. 633 DEGs). These changes were also qualitatively different and tissue specific with only ∼3% of the exercise training responsive changes shared between genotypes, (iWAT 32 DEGs, Liver 30 DEGs, Heart 33 DEGs and Lung 16 DEGs), (Figure 7B,E,H,K). Furthermore, predicted upstream transcription factor analysis of the exercise training responsive DEGs, similarly to skeletal muscle, showed largely different putative upstream regulators, Supplementary Figure 10. The magnitude and breadth of these differences were unexpected and point to the impact that training adaptations in skeletal muscle have on systemic adaptations to exercise.

**Figure 7:**
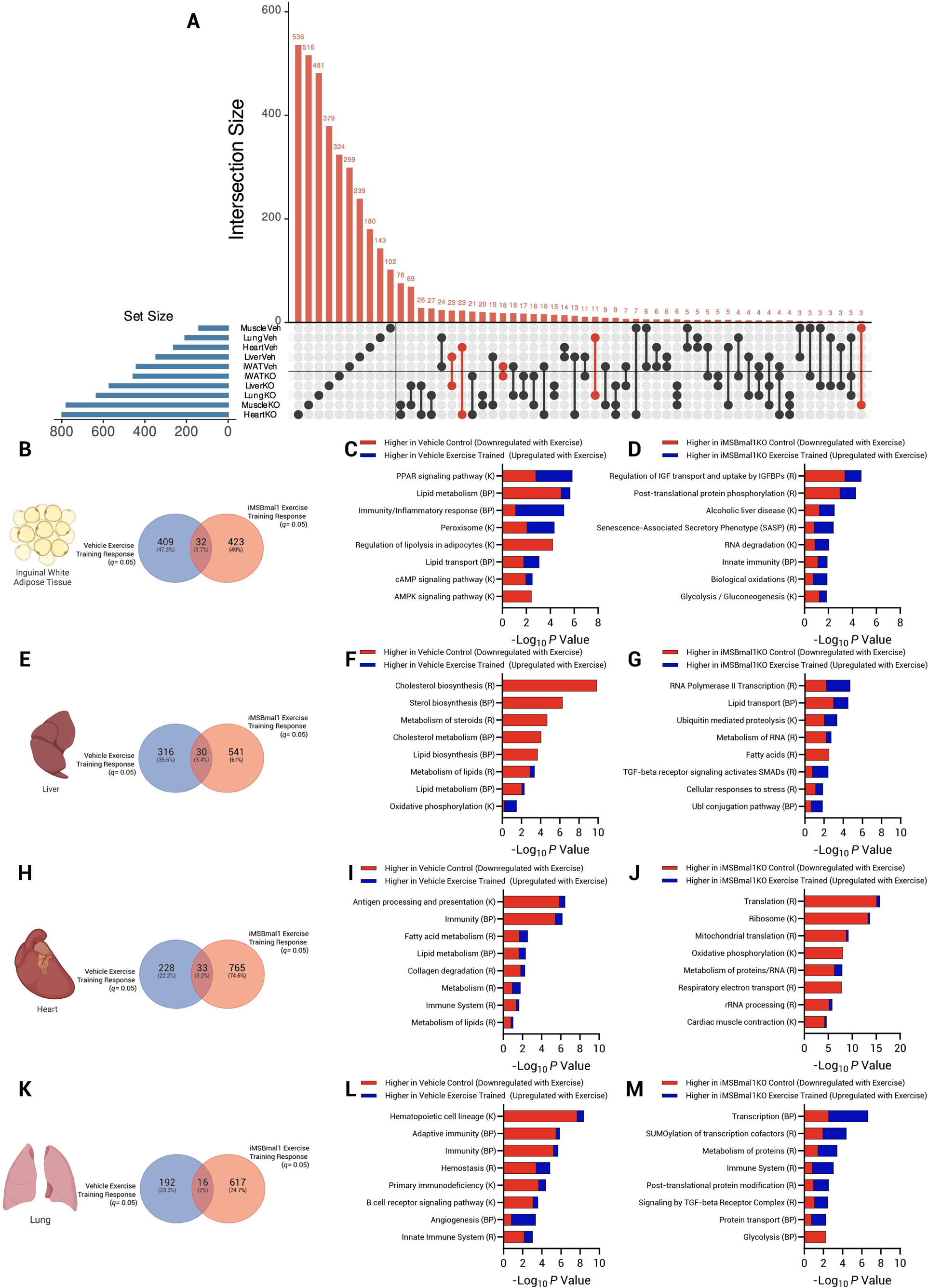
Divergent transcriptional reprogramming of peripheral tissues in response to endurance exercise training in the absence of skeletal muscle BMAL1. (A) Upset plot of the gene sets associated with the training response of each tissue in both genotypes. Set size indicates the total number of differentially expressed genes regulated by exercise training in each tissue in both genotypes. Numbers above bars indicate the number of differentially expressed genes. Overlapping differentially expressed genes which are common between tissues are indicated by the connected points below bars. Differentially expressed genes common between genotypes within the same tissue are indicated by red connected points. (B) Summary Venn diagram of genotype specific exercise training responsive genes in vehicle (409 DEGs) and iMSBmal1KO mice (423 DEGs) with shared features (32 DEGs) in inguinal white adipose tissue. Associated inguinal white adipose tissue pathway analysis for vehicle (C) and iMSBmal1KO mice (D). (E) Summary Venn diagram of genotype specific exercise training responsive genes in vehicle (316 DEGS) and iMSBmal1KO mice (541 DEGs) with shared features (30 DEGs) in liver tissue. Associated liver pathway analysis for vehicle (F) and iMSBmal1KO mice (G). (H) Summary Venn diagram of genotype specific exercise training responsive genes in vehicle (228 DEGs) and iMSBmal1KO mice (765 DEGs) with shared features (33 DEGs) in heart tissue. Associated heart pathway analysis for vehicle (I) and iMSBmal1KO mice (J). (K) Summary Venn diagram of genotype specific exercise training responsive genes in vehicle (192 DEGs) and iMSBmal1KO mice (617 DEGs) with shared features (16 DEGs) in lung tissue. Associated lung pathway analysis for vehicle (L) and iMSBmal1KO mice (M). Red segments of bars indicate downregulation with exercise training, whereas blue segments of bars indicate upregulation with exercise training. Associated pathway databases are indicated by (R) Reactome, (K) KEGG and (BP) Biological Processes.

In vehicle exercise trained iWAT, we observed an expected transcriptional remodeling with exercise training which included genes enriched in pathways associated with ‘PPAR signaling’, ‘Lipid Metabolism’ and ‘Inflammatory Response’. This included downregulation of *Pparg* itself along with other factors involved in lipid metabolism/storage, insulin sensitivity and inflammation. Analysis of the liver transcriptome of exercise trained vehicle mice demonstrated a synergy with the iWAT transcriptome, with downregulation of pathways including ‘Lipid/cholesterol biosynthesis/metabolism’, (Figure 7F). This common pathway enrichment between iWAT and liver was lost in the iMSBmal1KO training response. In iMSBmal1KO iWAT the exercise-responsive genes were associated with insulin resistant signatures and inflammation, included within ‘insulin-like growth factor transport/binding protein’ pathways. One example was the upregulation of the transcription factor *Srebf1*, known to promote cholesterol and fatty acid biosynthesis and downregulation of the glycolytic enzyme *Pkm*, (Figure 7C-D). Uniquely, liver of iMSBmal1KO exercise trained mice showed alterations in ‘RNA Polymerase II Transcription’, pathways associated with ubiquitin mediated proteolysis, and upregulation of a number of TGF-beta receptor signaling associated pathway genes including *Smad4*, *Smurf2*, *Itgav* and *Tgfbr1*, (Figure 7G). These results suggest that exercise training in the iMSBmal1KO mice leads to abnormal transcriptional remodeling in iWAT and liver including many biological signatures that could be considered ‘maladaptive’.

The exercise training response in the lung of the vehicle mice corroborated outcomes recently published investigations by the MoTrPAC consortium and were broadly related to decreased inflammation [50]. We identified 208 DEGs that are changed in the lung following training including 37 genes associated with the Reactome pathway ‘immune system’, over ∼83% of which were downregulated with exercise training. There were 633 training induced DEGs in the iMSBmal1KO lung which was a surprise as the transcriptomes of the sedentary vehicle and iMSBmal1KO lungs were not very different. This adaptation included 69 different ‘immune system’ associated genes were found to be enriched in the iMSBmal1KO exercise training response, 72% of which were upregulated with exercise training indicating a discordance in the inflammatory response with training between genotypes, (Figure 7K-M).

In the vehicle exercise trained heart we observed modest changes to the transcriptome associated with ‘Immunity’, ‘Lipid Metabolism’ and ‘Collagen Degradation’, (Figure 7H-I). Conversely, a majority of the exercise training responsive genes in the iMSBmal1KO heart were directionally opposite to the features identified in the loss of BMAL1 sedentary mice, see Figure 6D. This indicates that unlike the other peripheral tissues, exercise training rescued the dysregulation of these genes in the iMSBmal1KO heart, (Figure 6D/7J). This is supported by data presented in Supplementary Figure 11, where we highlight that 97% of genes in the heart that are disrupted by loss of muscle BMAL1 in sedentary mice, are exercise training responsive in iMSBmal1KO hearts and return closer to vehicle control expression levels, R^2^=0.67. The mechanism(s) through which exercise training restores the heart transcriptome in the iMSBmal1KO mouse is unclear and requires additional study.

### 3.9. Exercise training in the abence of skeletal muscle BMAL1 alters the steady state immune system and metabolic pathways of peripheral tissues

Since inflammation and metabolism pathways were the most common differences among the transcriptional changes, we computed Z-scores for each pathway by averaging the Z-score of every gene found within each pathway. Specifically, we included ‘Immune System’, ‘Carbohydrate Metabolism’, ‘Lipid Metabolism’, ‘Amino Acid Metabolism’ and the ‘Endocrine System’ to characterize between genotype changes in the exercise trained state. As visualized in Figure 8A, genes associated with the ‘Immune System’ pathways were generally expressed at a higher level with exercise training in skeletal muscle and iWAT and decreased in liver heart and lung in the vehicle mice, a signature that has been linked to increased immune cell recruitment [50]. In contrast, this pattern is reversed in the iMSBmal1KO tissues following training with expression of these immune pathway genes lower in the muscle and iWAT and much higher in liver, lungs and heart with training. These observations indicate that the altered tissue environment(s) and/or altered immune cells in circulation results in recruitment or retention of different amounts, or types of immune cell populations in these tissue and organs with targeted loss of muscle BMAL1 and warrants further investigation.

**Figure 8:**
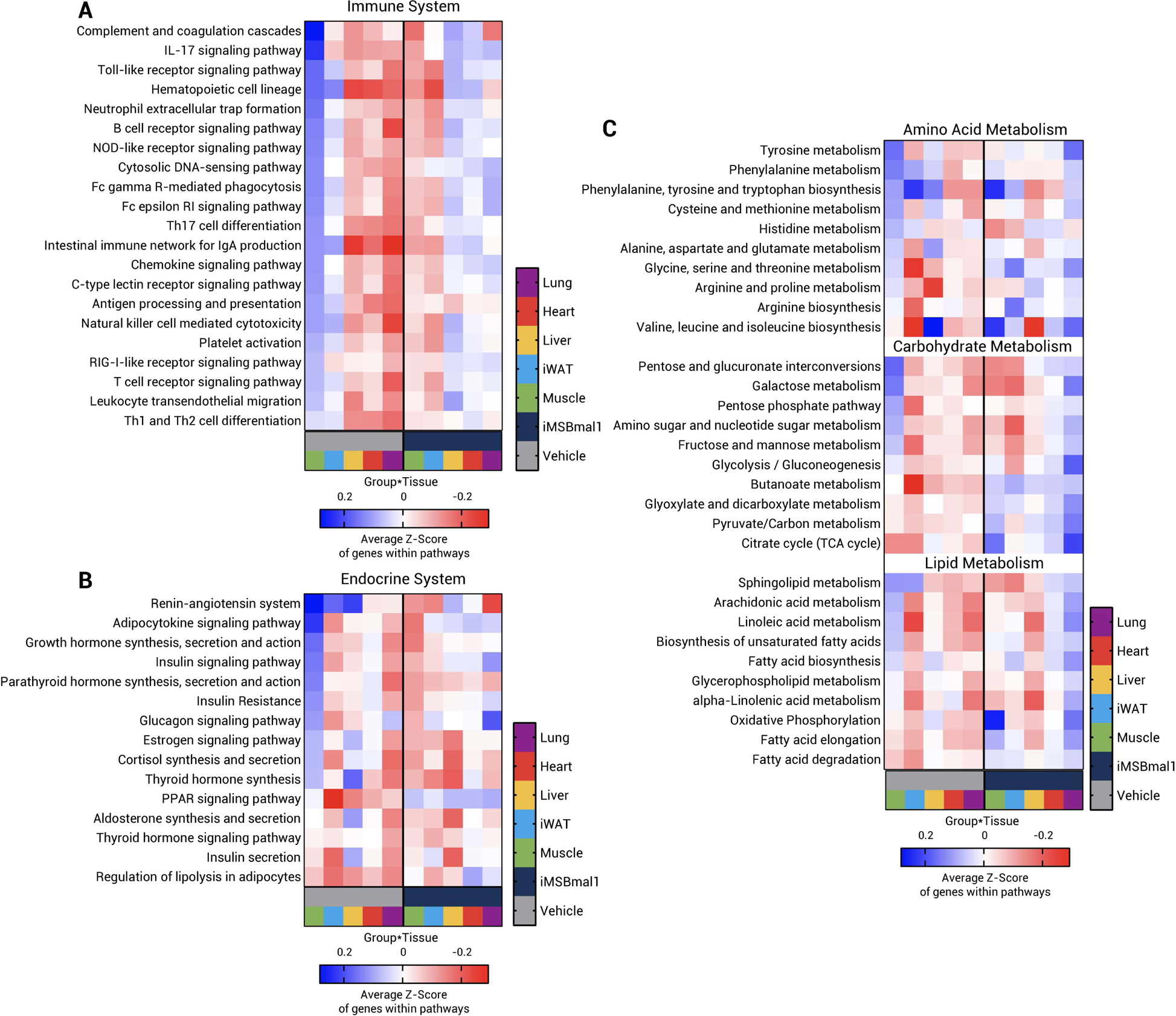
Loss of skeletal muscle BMAL1 causes rewiring of inflammatory and metabolic pathways across multiple tissues after exercise training. Deseq2 normalized counts were used to produce within tissue Z-scores for all expressed genes for all 4 experimental groups. Average Z-scores were then produced from all genes found within the given KEGG pathway for vehicle exercise trained (grey bars) and iMSBmal1KO exercise trained mice (dark blue bars) for each tissue. Tissues are indicated by green bars (Gastrocnemius Muscle), light blue bars (iWAT), yellow bars (Liver), red bars (Heart) and purple bars (Lung). Data is presented for KEGG pathways related to (A) the Immune System, (B) Endocrine System and (C) Amino Acid, Carbohydrate and Lipid Metabolism. Blue signifies relative upregulation of associated pathways and red signifies relative downregulation of the genes within the associated pathways.

The endocrine system pathways were also differentially modified by exercise training between genotypes. One recognized exercise feature is the ‘PPAR signaling pathway’ which was signficiantly lower across tissues in the vehicle mice but was higher across all 5 tissues in the iMSBmal1KO exercise trained mice, Figure 8B. This likely reflects the altered energy homeostasis resulting from loss of skeletal muscle BMAL1 promoting systemic increases in fatty acid metabolism [60–62]. As expected from our muscle transcriptome and metabolomic data, we observed strong upregulation of pathways associated with amino acid metabolism that are generally expressed at a higher levels in all peripheral tissues in the iMSBmal1KO exercise trained mice, compared with their vehicle trained counterparts, Figure 8C. In particular, we identify the lung as a tissue that gets brought into substrate metabolism in a much bigger way with exercise in the iMSBmal1KO mouse. This is a unique observation that highlights the potential for lung metabolism to contribute to systemic energy homeostasis with exercise training. Together, analysis of these pathways emphasise that skeletal muscle BMAL1, and likely a functional skeletal muscle clock, is essential for systemic tissue homeostasis and peripheral tissue transcriptional remodelling in response to exercise training, pertaining to metabolic and immune components and these warrant further investigation.

## 4. Discussion

In this investigation, we addressed the contribution of the core circadian clock factor, BMAL1, in skeletal muscle to both acute transcriptional responses to exercise and transcriptional remodelling in response to exercise training. Additionally, we adopted a systems biology approach to investigate how loss of skeletal muscle BMAL1 altered peripheral tissue homeostasis as well as exercise training adaptations in iWAT, liver, heart, and lung of male mice.

Mice deficient in muscle BMAL1 failed to mount a ‘normal’ transcriptional response to acute exercise despite performing the same amount of treadmill work. iMSBmal1KO mice demonstrated a blunted transcriptional response to acute exercise and this was associated with the lack of up-regulation of *Ppargc1a*, *Fos*, and orphan nuclear receptors, *Nr4a1* and *Nr4a3*. This was concomitant with the lack of activation of genes within well-known exercise responsive pathways, e.g., PI3k-Akt, MAPK and PPAR. We also identified that signal transduction associated genes were significantly altered at both the chromatin accessibility level and transcript level, consistent with reports in liver[63], that BMAL1 plays an important role in epigenetic regulation as well as transcriptional regulation.

We next tested the exercise training response in the mice lacking BMAL1 specifically in skeletal muscle. As with the acute response studies, the mice were able to perform the same training paradigm so that the treadmill work for both vehicle and iMSBmal1KO mice was the same over a 6-week period. Analysis of the transcriptome identified a novel BMAL1-dependent program of gene expression in skeletal muscle in response to chronic exercise training. Despite the same exercise training volumes, the adaptation of the transcriptome in skeletal muscle resulted in a quantitatively larger reprogramming with very few genes in common (∼3%), representing very different enriched pathways and putative upstream transcriptional networks. It is important to note that of the subset of common exercise responsive genes between the two genotypes (*n*=63), only half of them change in the same direction. These outcomes indicate that the well-defined molecular responses to a set exercise training paradigm in skeletal muscle requires BMAL1 and/or a functional circadian clock mechanism, but in its absence requires a far larger network of genes to elicit adaptations. We also note that although the transcriptome was largely divergent between genotypes, classic markers of muscle, such as fiber size and fiber type, were not different indicating that these parameters of muscle are not under BMAL1 control.

Our multi-omic analyses identified that the skeletal muscle without BMAL1 did adapt but the steady state responses within the transcriptome and metabolome were quantitatively larger and importantly, fundamentally divergent. This included the observation that exercise training is not sufficient to rescue the dysregulation of genes essential for glucose metabolism in the iMSBmal1KO skeletal muscle demonstrating that exercise training is not sufficient to restore metabolic flexibility in the absence of BMAL1. To support the increased energy demand of exercise training the iMSBmal1KO muscle exhibited a uniform up-regulation of genes, likely regulated by *Esrra*, to support the TCA cycle, beta oxidation and electron transport complexes. These divergent transcriptomic changes in the iMSBmal1KO muscle were supported by the divergent metabolomic responses to training, with amino acids being an important source for TCA substrates to support oxidative metabolism. These studies uncover that in response to the increased energy demand with exercise the iMSBmal1KO muscle that adaptation can occur, but it does not resemble the expected molecular and metabolic adaptations seen in the vehicle treated mice.

Our findings have similarities to recent work demonstrating the contribution of muscle specific *Ppargc1a* to the exercise training response [42]. This is not surprising as it has been shown previously that *Ppargc1a* is a circadian clock output gene in skeletal muscle. In support of this, our acute exercise studies extend this relationship as a clock output gene and demonstrate that up-regulation of *Ppargc1a* requires muscle BMAL1. This places muscle BMAL1, and likely the core clock mechanism, as an upstream regulator of *Ppargc1a* expression in response to exercise. Recent work has demonstrated similar relationships between the core clock and time-of-day exercise activation of HIF1A/HIF1A target genes [17]. We suggest that this loss of the exercise expression of *Ppargc1a* and the other acute exercise transcription factors are significant contributors to the divergent exercise training transcriptome in the iMSBmal1KO mouse. We found that only 3% of the differentially expressed genes were shared between vehicle treated and iMSBmal1KO muscle. This is a significant reduction compared to the 32% shared differentially expressed genes defined following training in the muscle specific *Ppargc1a* KO mice. Thus, our findings establish the importance of muscle BMAL1 as an upstream regulator of *Ppargc1a* and other transcription factors, including *Nr4a3*. We suggest that the loss of the ability of exercise to induce these transcription factors result in a largely divergent molecular adaptation to exercise training [42].

There is growing recognition that exercise is a lifestyle intervention that contributes to systemic health with beneficial impacts on virtually all organs studied to date. While this is commonly recognized, the molecular mechanisms for these broad adaptations are still poorly understood. A key finding from this work is that we have identified that muscle specific adaptations with exercise are significant contributors to molecular adaptations in peripheral tissues. We were surprised to find that peripheral tissue remodelling, similarly, to skeletal muscle was greater in magnitude in every tissue in the iMSBmal1KO mouse. The changes in gene expression were largely unique to each tissue with <4% overlap in the exercise training responsive genes across iWAT, liver, heart, and lung between the vehicle and iMSBmal1KO mice. Importantly, while exercise training is known to be beneficial to the resilience of most tissues, the analysis of the functional groups of genes that change with training in the iMSBmal1KO mice suggest several organs, including lungs, liver and iWAT may be exhibiting a maladaptive response. Since the loss of BMAL1 only occurred in skeletal muscle these outcomes highlight the systemic tissue cross talk induced by skeletal muscle in response to exercise training. We speculate that these systemic changes are a result of two factors, or the combination of both. One factor is the systemic metabolic changes which result from the metabolic inflexibility of the BMAL1 deficient skeletal muscles exacerbated by the energetic demands of exercise training. Changes in metabolite/substrate availability/abundance may subsequently influence the transcriptomic state of the cells within the peripheral tissues. Secondly, our data indicate that loss of skeletal muscle BMAL1 changes the inflammatory transcriptome both within the muscle and as well as peripheral tissues, and this altered status may interfere with the adaptive processes within these tissues. Our work adds further evidence to the growing body of literature demonstrating the importance of BMAL1 in maintaining homeostasis and facilitating adaptive processes in different and across organ systems [64–70].

In conclusion, our investigation has uncovered the critical role that BMAL1 plays in skeletal muscle as a key regulator of the gene expression programs for both acute exercise and training adaptations. In addition, our work has uncovered the significant impact that altered exercise response in muscle plays in the peripheral tissue adaptation to exercise training. Specifically, we propose that muscle BMAL1 acts upstream and is necessary for the well-defined acute exercise induction of key transcription factors including *Ppargc1a*. We also note that the transcriptome adaptations to steady state training suggest that without BMAL1 the muscle does not achieve the expected homeostatic program. Lastly, we propose that the peripheral tissue adaptations to exercise training are tightly coupled to the adaptations in skeletal muscle. In fact, if the muscle adaptations diverge to a more maladaptive state this is also linked to increased inflammation across many tissues. Understanding the molecular targets and pathways contributing to health vs. maladaptive exercise adaptations will be critical for the next stage of therapeutic design for exercise mimetics.

## Acknowledgements

We would like to thank Dr. Daniel Kopinke at the University of Florida for the use of his microscope. We also extend our thanks to the Gene Expression & Genotyping (GE) core, RRID:SCR_019145, and NextGen DNA Sequencing (NS) core, RRID:SCR_019152 and the Southeast Center for Integrated Metabolomics, (SECIM) at the University of Florida for library preparation, DNA sequencing and metabolomics respectively. We would like to thank Dr. Jerome Menet for constructive feedback and discussions on our data. This work was supported by NIH R01AR079220 to KAE.

## Author contributions

M.R.V, S.J.H, K.A.E conceived and designed research. M.R.V, S.J.H, M.A.G, C.A.W, R.A.M, H.E.B, I.G.J performed experiments. M.R.V, M.A.G, Z.H analysed data. M.R.V, R.A.M, M.A.G, K.A.E interpreted results of experiments. M.R.V prepared figures, M.R.V, K.A.E drafted manuscript. M.R.V, K.A.E edited and revised manuscript, M.R.V, S.J.H, M.A.G, C.A.W, R.A.M, H.E.B, I.G.J, Z.H, K.A.E approved final version of manuscript.

## Supplemental Files

Supplementary File 1: RNA/ATAC-seq Data

Supplementary File 2: LCMS Metabolomics Data

## Supplemental Figure Legends

**Supplementary Figure 1:**
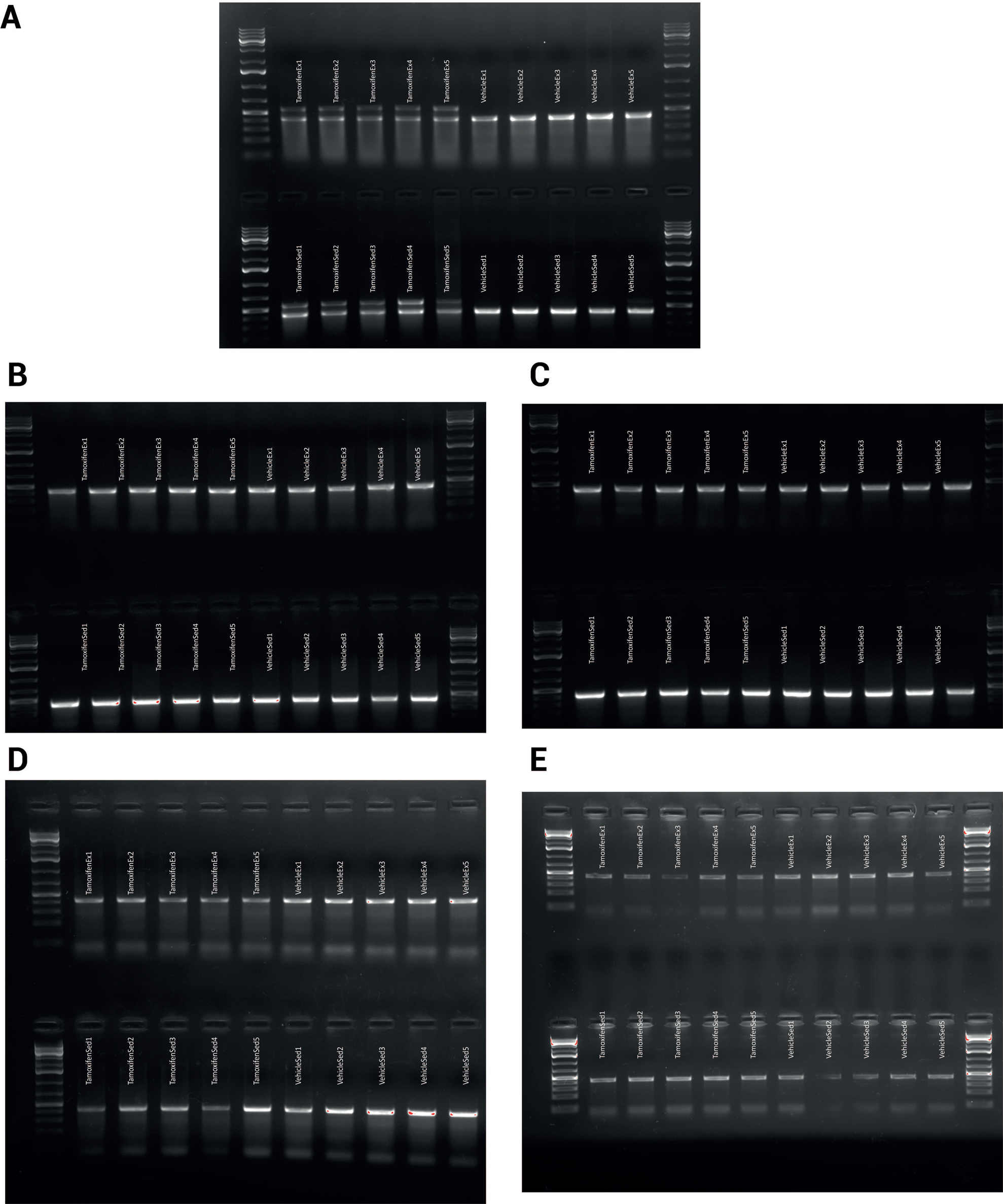
(A) Confirmation of successful Bmal1 Cre-Recombination in gastrocnemius muscles and lack of recombination in heart (B), liver (C), inguinal white adipose tissue (D) and lung (E).

**Supplementary Figure 2:**
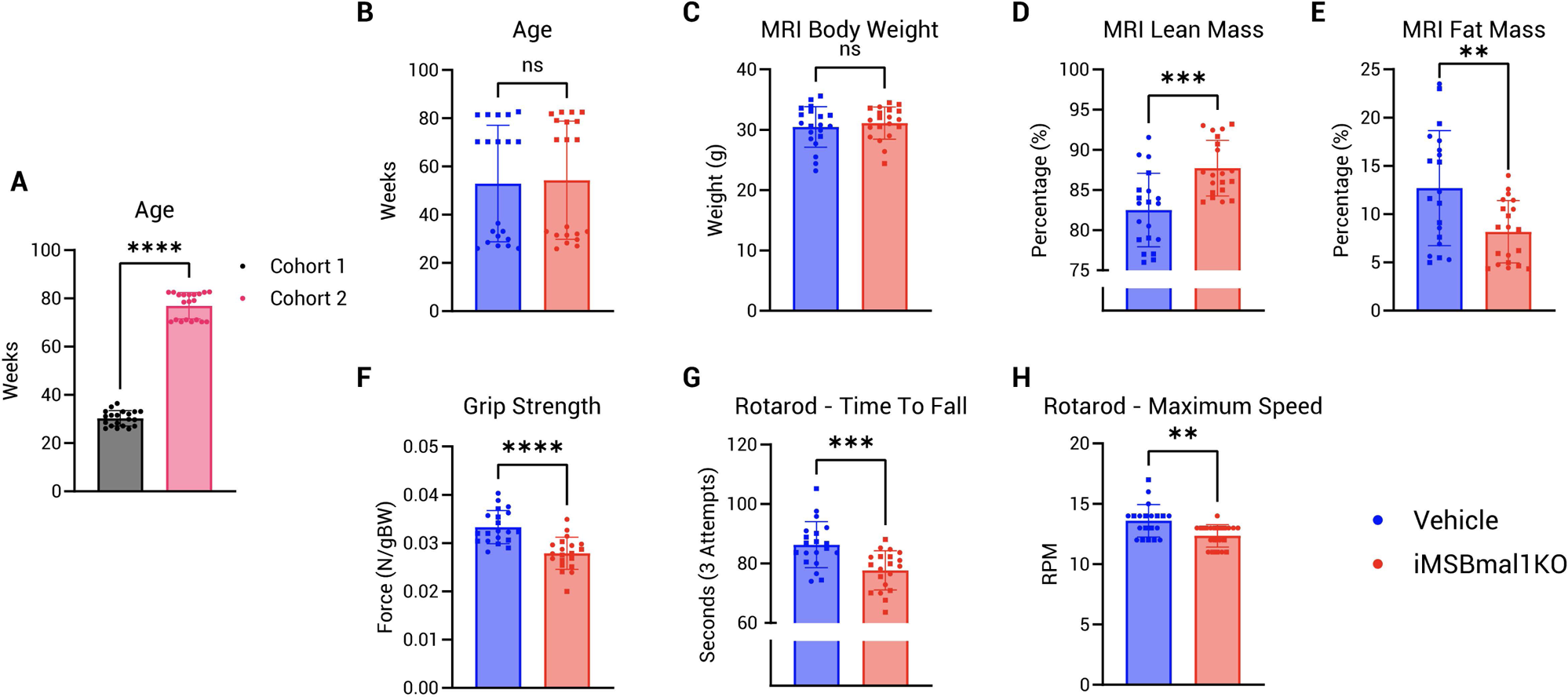
Physiological assessment of vehicle treated and iMSBmal1KO mice. (A) Differences in age (weeks) between cohorts 1 and 2 at termination. Mice in the older cohort are indicated by square data points, mice in the younger cohort are indicated by circle data points. (B) Age in weeks (termination). (C) Body Weight (g). (D) MRI measured lean mass as a percentage of total body weight. (E) MRI measured fat mass as a percentage of total body weight. (F) Forelimb grip strength (N/gBW). Rotarod time to fall assessment in seconds (average of 3 trials) (G) and maximal speed (RPM) (H). Cohort 1 is delineated by circles, cohort 2 is delineated by squares. Vehicle treated mice are delineated by blue circles, iMSBmal1KO mice are delineated by red circles. Significance was determined by use of an unpaired t-test where ns indicates *p*>0.05, * indicates *p*<0.05, ** indicates *p*<0.01, *** indicates *p*<0.001 and **** indicates *p*<0.0001.

**Supplementary Figure 3:**
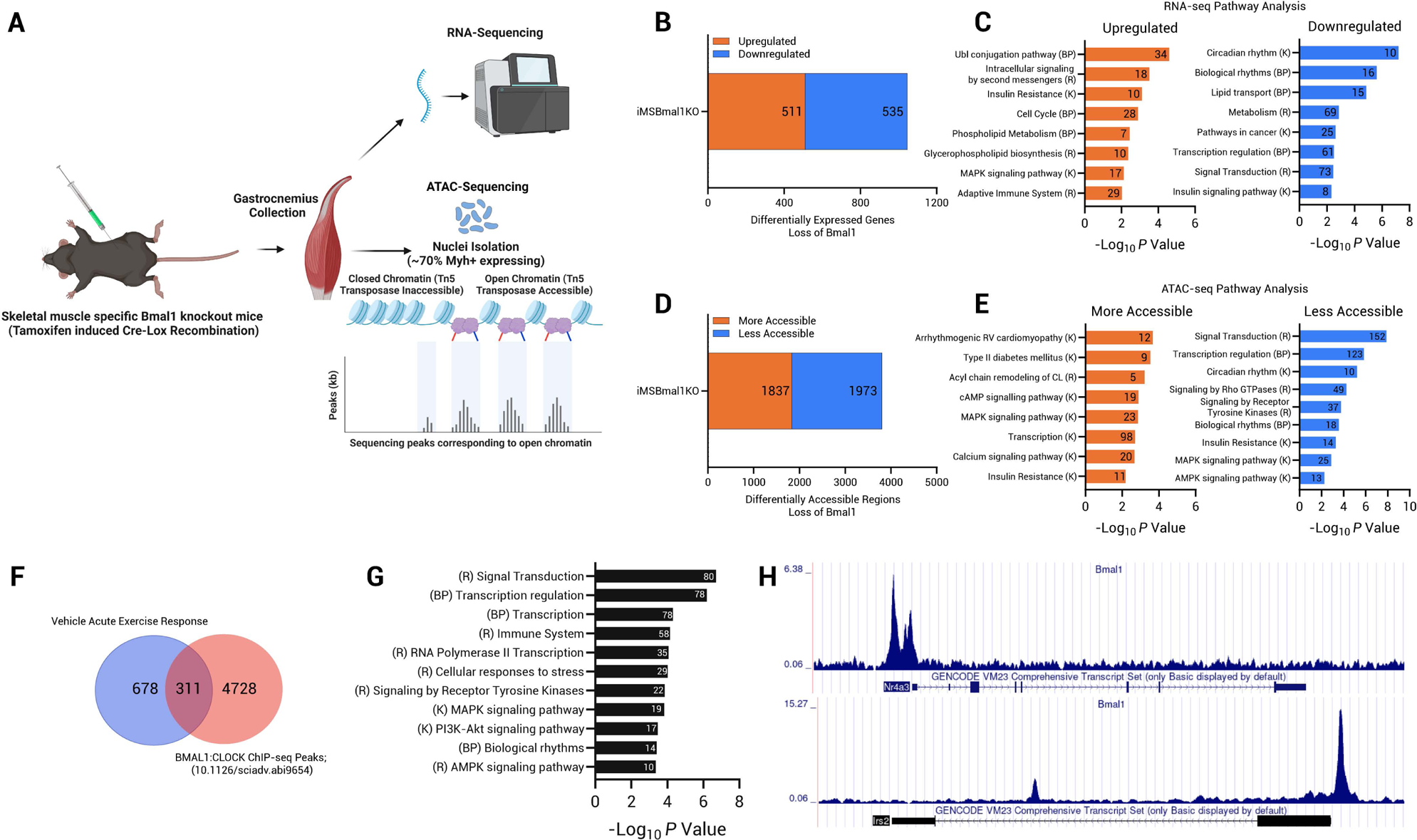
Transcriptome and chromatin accessibility analysis of iMSBmal1KO mice. Vehicle treated and iMSBmal1KO mice underwent baseline physiological phenotyping including RNA and ATAC-sequencing analysis at ZT13 on gastrocnemius tissue. (B) Number of DEGs genes with loss of skeletal muscle *Bmal1* (*q* = 0.05). Numbers in bars indicate up or downregulation. (C) Pathway analysis of differentially expressed RNAs split by direction of change. Numbers in bars indicate the number of genes present within the pathway. (D) Number of differentially accessible regions as assessed by ATAC-seq. Numbers in bars indicate the number of DNA regions/peaks becoming more or less accessible with loss of *Bmal1*. (E) Pathway analysis of differentially accessible chromatin regions split by direction of change. Numbers in bars indicate the number of genes present within the pathway. (F) Venn overlap of vehicle acute exercise response DEGs and genes regulated by skeletal muscle BMAL1 (BMAL1:CLOCK ChIP-seq peaks, Gabriel *et al*. 2021 Sci Adv). (G) Pathway analysis of overlapping features. (H) Example BMAL1 ChIP-seq plots illustrating BMAL1 binding in promoter regions of *Nr4a3* and *Irs2*.

**Supplementary Figure 4:**
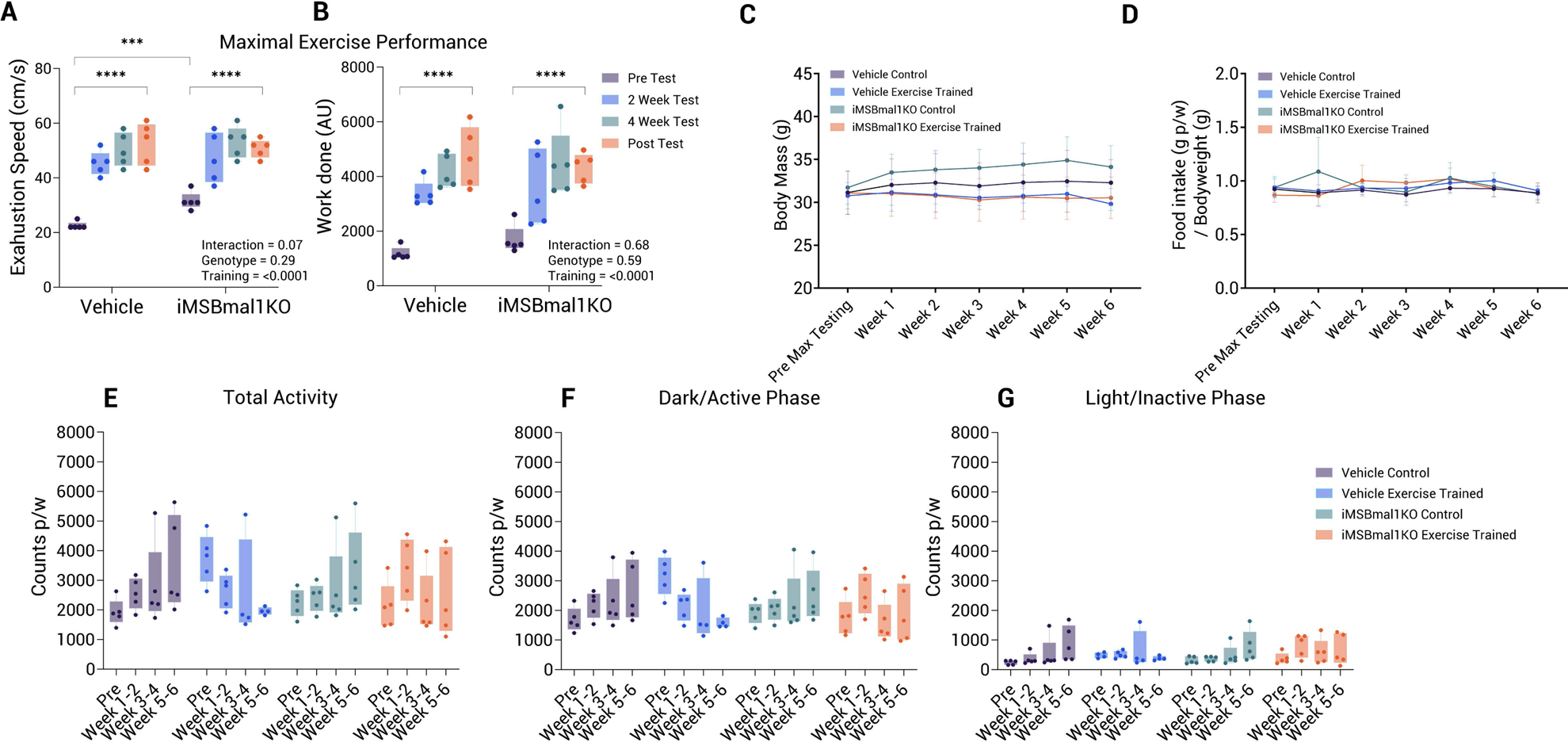
Panel (A/B) Maximum work-done tests were performed prior to, after 2, 4 and 6-weeks of training with the speed adjusted accordingly to make training progressive. (C) The amount of food consumed per week was measured and is presented as a proportion of body weight. (D) Measurements of body mass were taken weekly. Habitual cage activity as measured by infrared cage monitors revealed no change in total activity levels between groups. (E) Total Activity. (F) Activity restricted to the dark/active phase. (G) Activity restricted to the light/inactive phase. Significance for two-way ANOVA analysis is presented as \**p* ≤ 0.05, \*\**p* ≤ 0.01, \*\*\**p* ≤ 0.001, \*\*\*\**p* ≤ 0.0001.

**Supplementary Figure 5:**
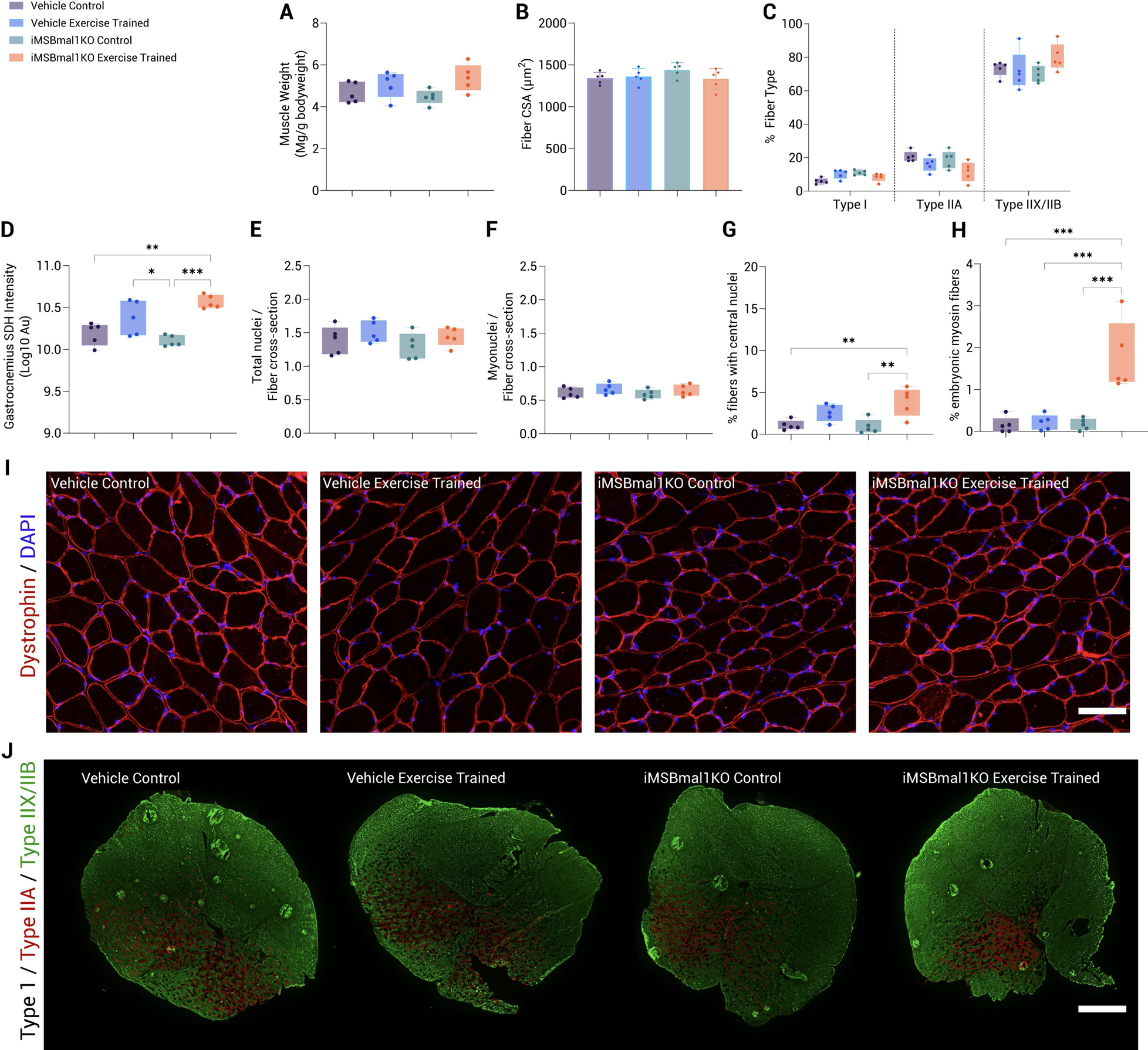
Histological phenotyping of gastrocnemius muscle. (A) Gastrocnemius muscle wet weight is presented as milligrams per gram of body weight. Myovision 2.0 software was used to automatically assess the average fiber cross-sectional area (µm2) (B), the proportion of different fiber types (expressed as a percentage of all fibers) (C). (D) Gastrocnemius SDH intensity value (Log10 AU). Myovision 2.0 assessments of the total number of nuclei per fiber cross-section (E), the number of myonuclei per fiber cross-section (F). (G) illustrates the % of fibers with central nuclei and (H) illustrates the % of fibers that were embryonic myosin positive. (I) Representative dystrophin and DAPI images for each respective group. Scale bars indicate 100µm. (J) Representative immunofluorescent images depicting whole cross-section gastrocnemius muscles from each experimental condition labeled with Type IIA/Type IIX/B. Scale bars indicate 1800µm. Data are presented as mean ± standard deviation or min-max values for box plots. Significance for one-way ANOVA analysis is presented as \**P* ≤ 0.05, \*\**P* ≤ 0.01, \*\*\**P* ≤ 0.001.

**Supplementary Figure 6:**
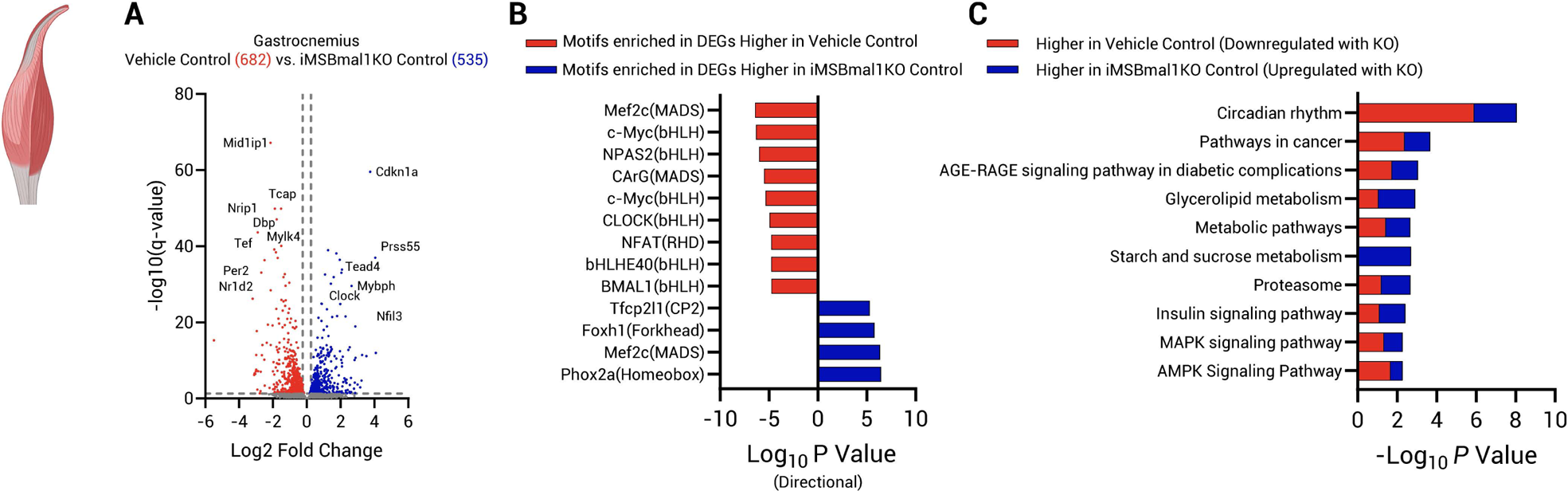
Training Study. Loss of Bmal1 in skeletal muscle causes transcriptional disruption of circadian rhythms, substrate metabolism and energy sensing pathways. (A) Volcano plot illustrating the DEGs between vehicle control and iMSBmal1KO controls. A total of 1217 (682 downregulated, 535 upregulated) genes were differentially expressed using an FDR cut off of <0.05. (B) HOMER Motif analysis performed on up and downregulated DEGs (*q*=0.05), identifies a number of transcription factors regulating the transcriptional response to loss of BMAL11 in skeletal muscle. (C) KEGG Pathway analysis reveals disruption of circadian rhythms, metabolic pathways and cell cycle/energy sensing pathways with loss of BMAL1 in skeletal muscle.

**Supplementary Figure 7:**
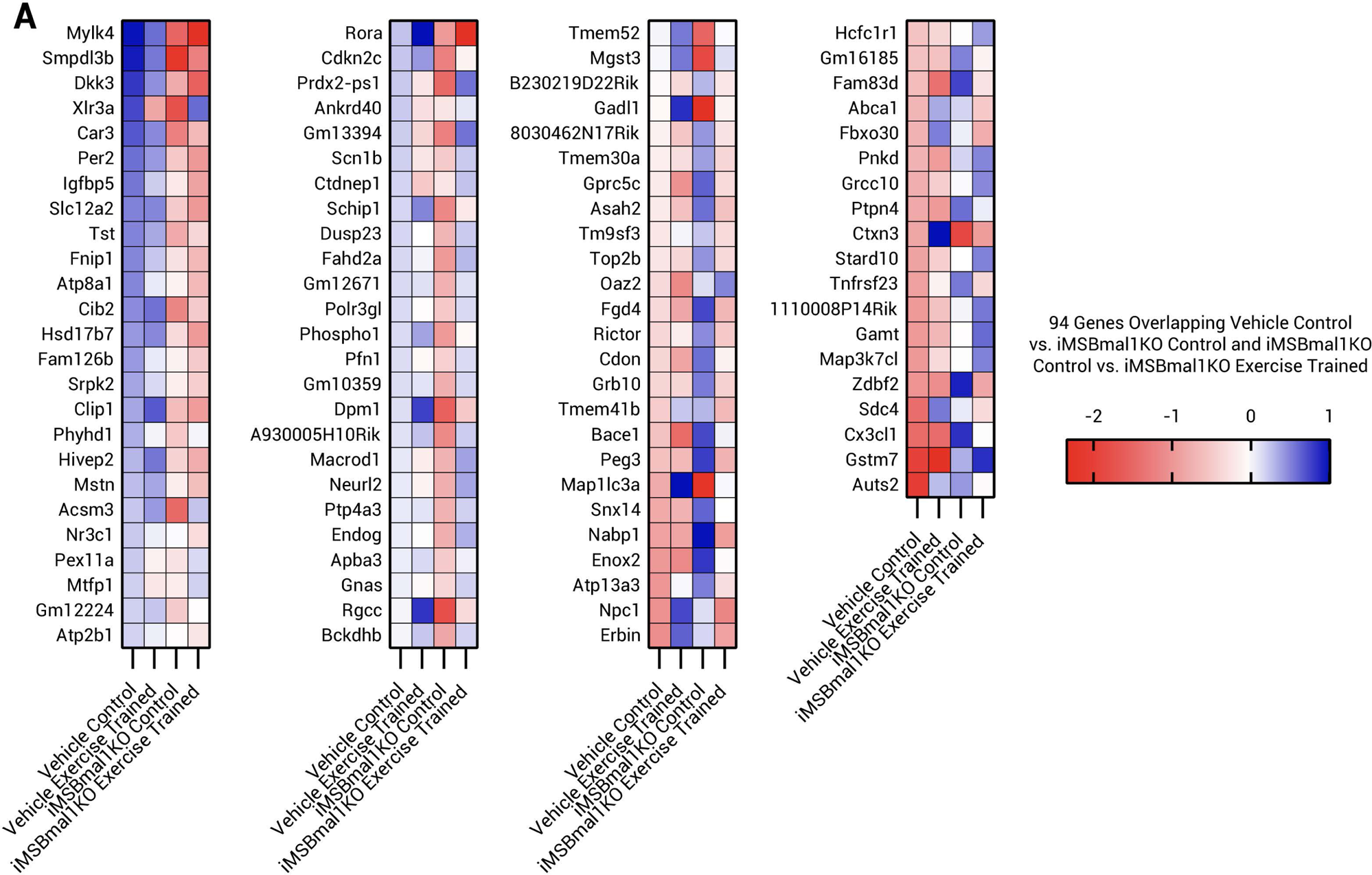
(A) Heatmap representing the 94 genes overlapping the vehicle control vs. iMSBmal1KO Control statistical comparison and iMSBmal1KO Control vs. iMSBmal1KO exercise trained statistical comparison.

**Supplementary Figure 8:**
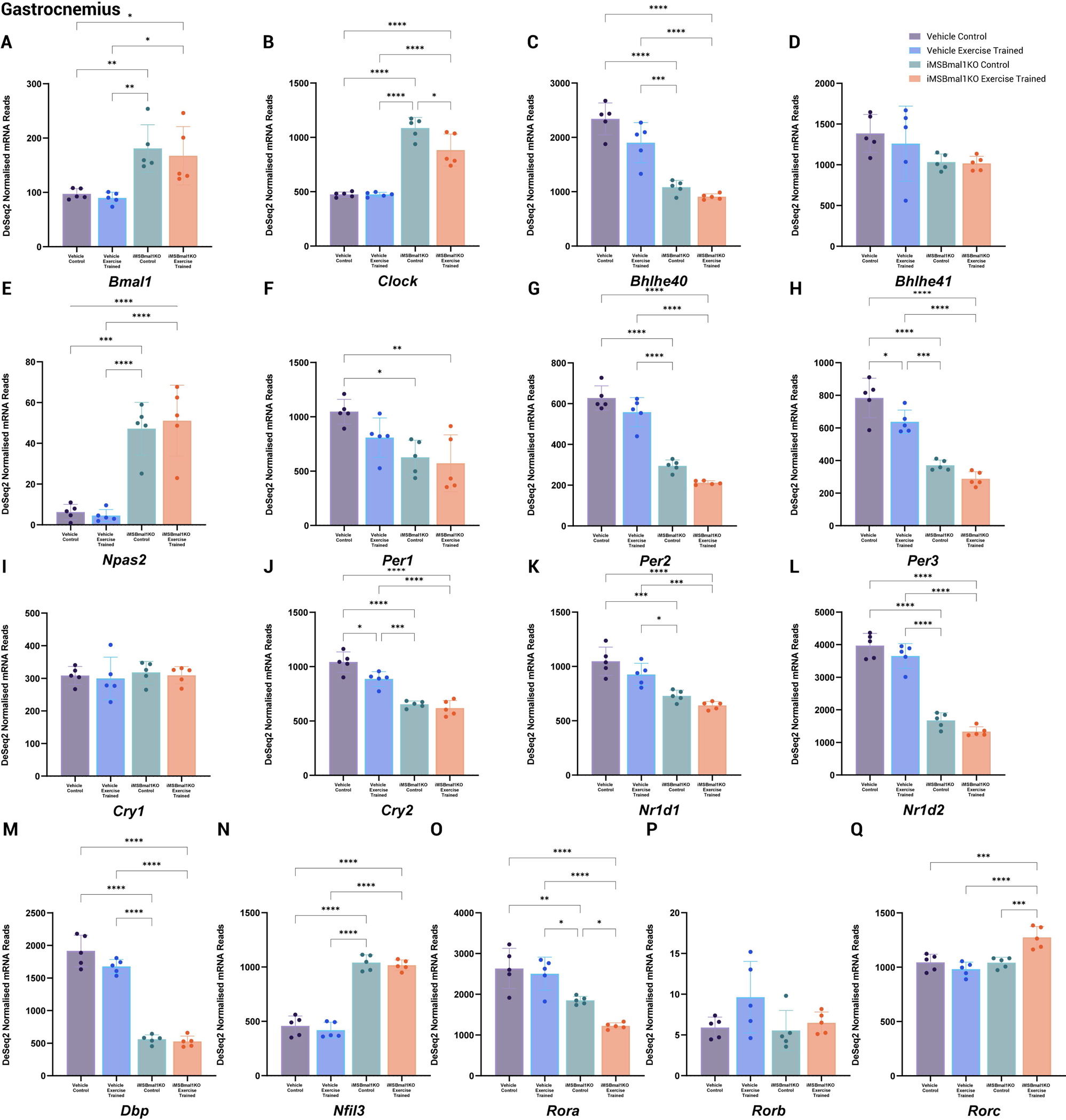
ZT13 core clock factor gene expression (Deseq2 normalized counts) in the gastrocnemius muscle in response to muscle specific Bmal1KO, exercise training and the combination of the two treatments. Data is presented as mean ± standard deviation. Significance for one way ANOVA analysis is presented as \**P* ≤ 0.05, \*\**P* ≤ 0.01, \*\*\**P* ≤ 0.001, \*\*\*\**P* ≤ 0.0001.

**Supplementary Figure 9:**
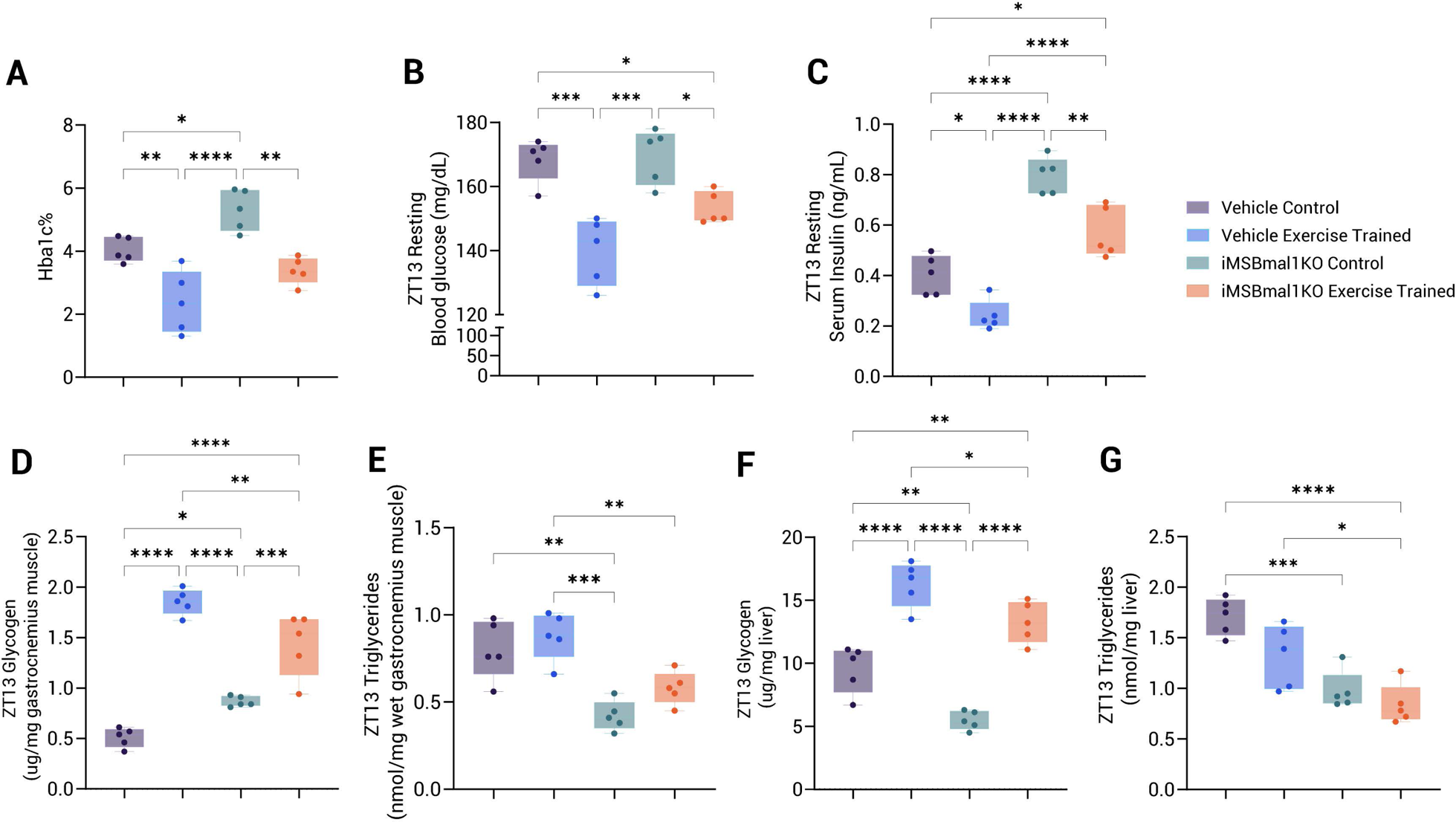
Metabolic indices of vehicle treated and iMSBmal1KO mice with and without 6-weeks exercise training taken at ZT13. (A) Blood hemoglobin A1C percentage (Hba1c%), (B) resting blood glucose (mg/dL), (C) resting plasma insulin (ng/dL), (D) gastrocnemius muscle glycogen concentration (µg/mg), (E) gastrocnemius muscle triglyceride concentration (nmol/mg), (F) liver glycogen concentration (µg/mg), (G) liver triglyceride concentration (nmol/mg). Significance for one way ANOVA analysis is presented as \**p* ≤ 0.05, \*\**p* ≤ 0.01, \*\*\**p* ≤ 0.001, \*\*\*\**p* ≤ 0.0001.

**Supplementary Figure 10:**
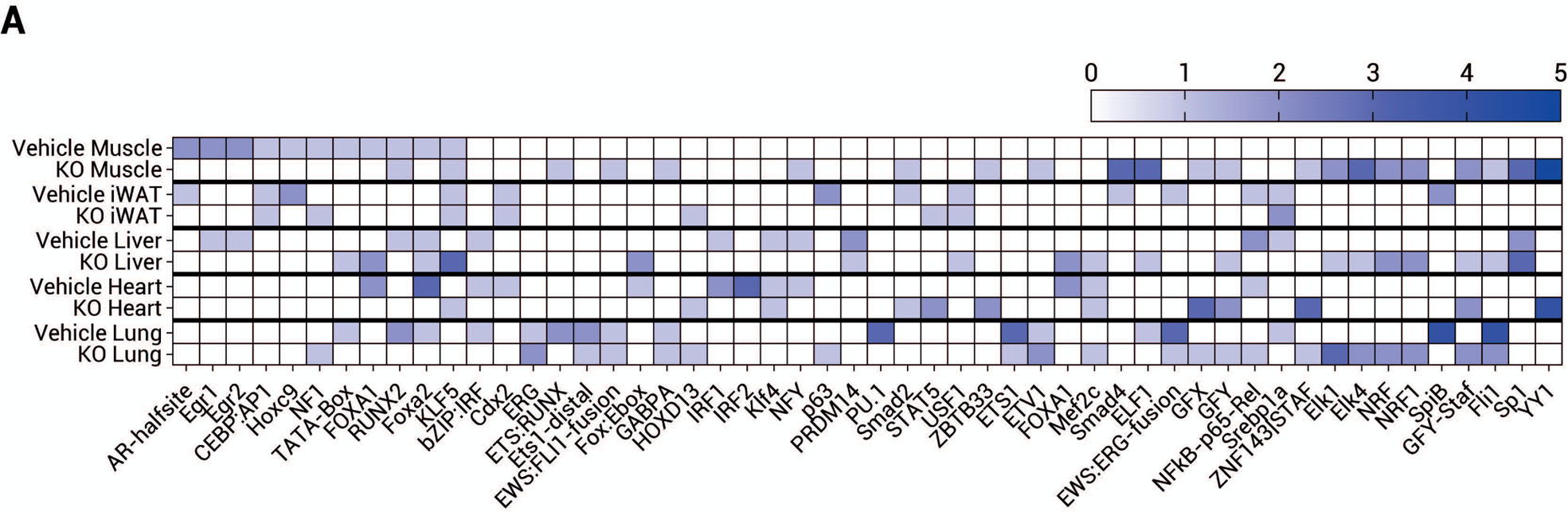
(A) HOMER transcription factor enrichment analysis of exercise responsive DEGs across genotypes and tissues presented as −Log10 *P*-values.

**Supplementary Figure 11:**
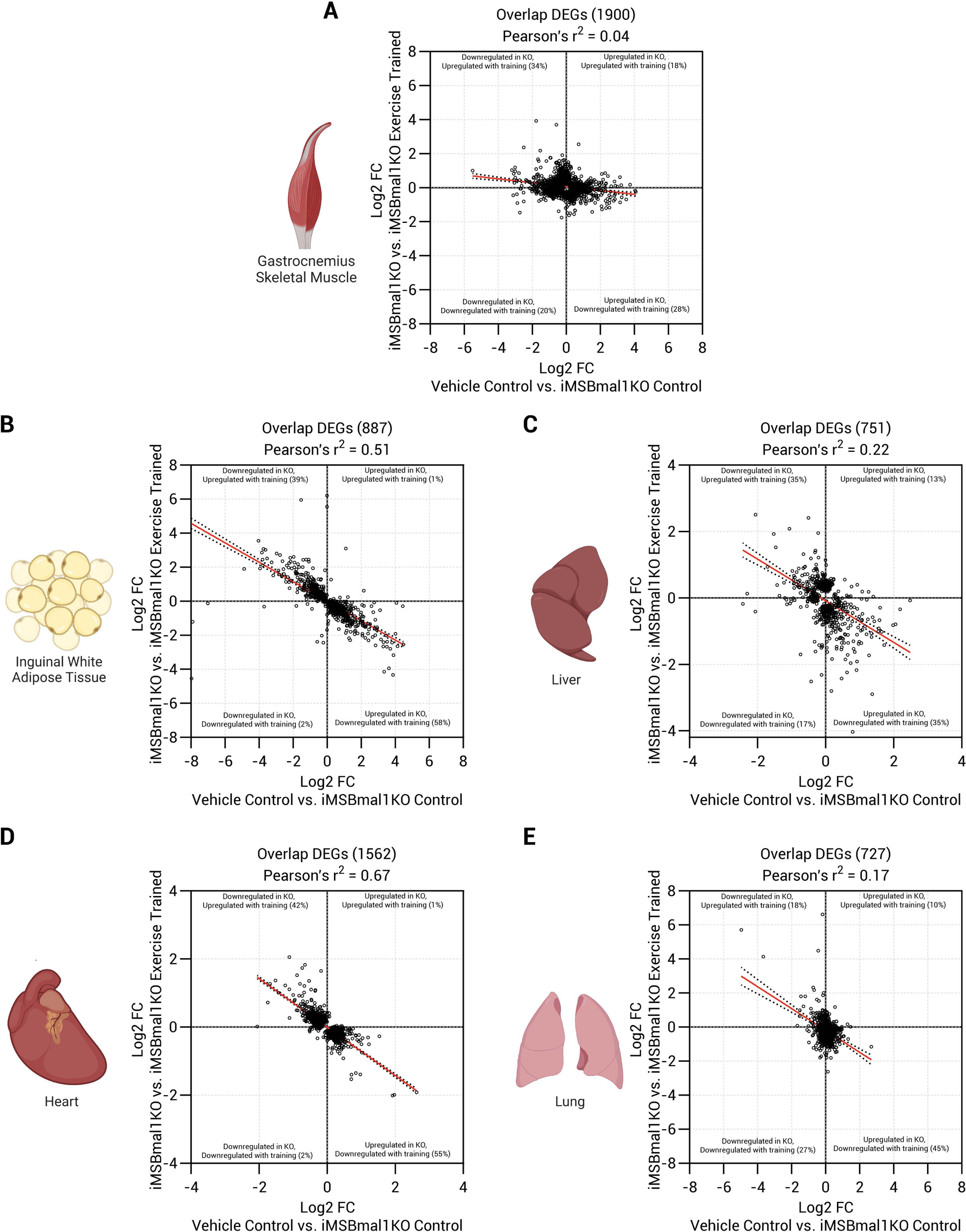
Rescue correlations for all genes affected by loss of skeletal muscle BMAL1 and their exercise trained responses in iMSBmal1KO mice for (A) gastrocnemius muscle, (B) inguinal white adipose tissue, (C) liver, (D) heart and (E) lung. iWAT and heart display relatively strong relationships (R^2^= 0.51-0.67) suggesting exercise training in the iMSBma1KO mice brings their transcriptomes closer to vehicle control mice. Muscle, liver and lung show weaker relationships (R^2^= 0.04-0.22).

