## Supplementary Figures for "Skeletal muscle BMAL1 is necessary for transcriptional adaptation of local and peripheral tissues in response to endurance exercise training"

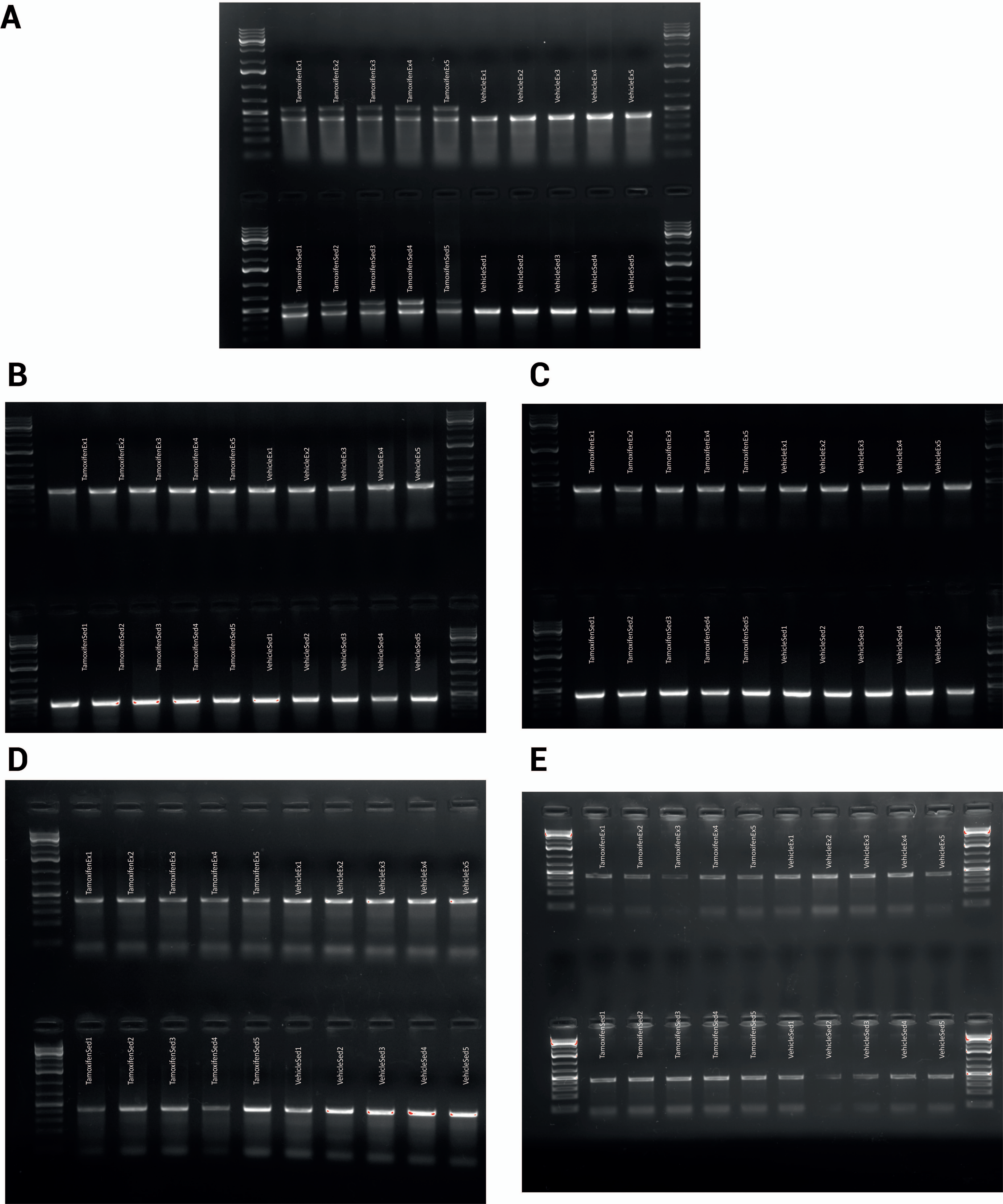
A
TamoxifenEx1
TamoxifenEx2
TamoxifenEx3
TamoxifenEx4
TamoxifenEx5
VehicleEx1
VehicleEx2
VehicleEx3
VehicleEx4
VehicleEx5
TamoxifenSed1
TamoxifenSed2
TamoxifenSed3
TamoxifenSed4
TamoxifenSed5
VehicleSed1
VehicleSed2
VehicleSed3
VehicleSed4
VehicleSed5
B
C
D
E

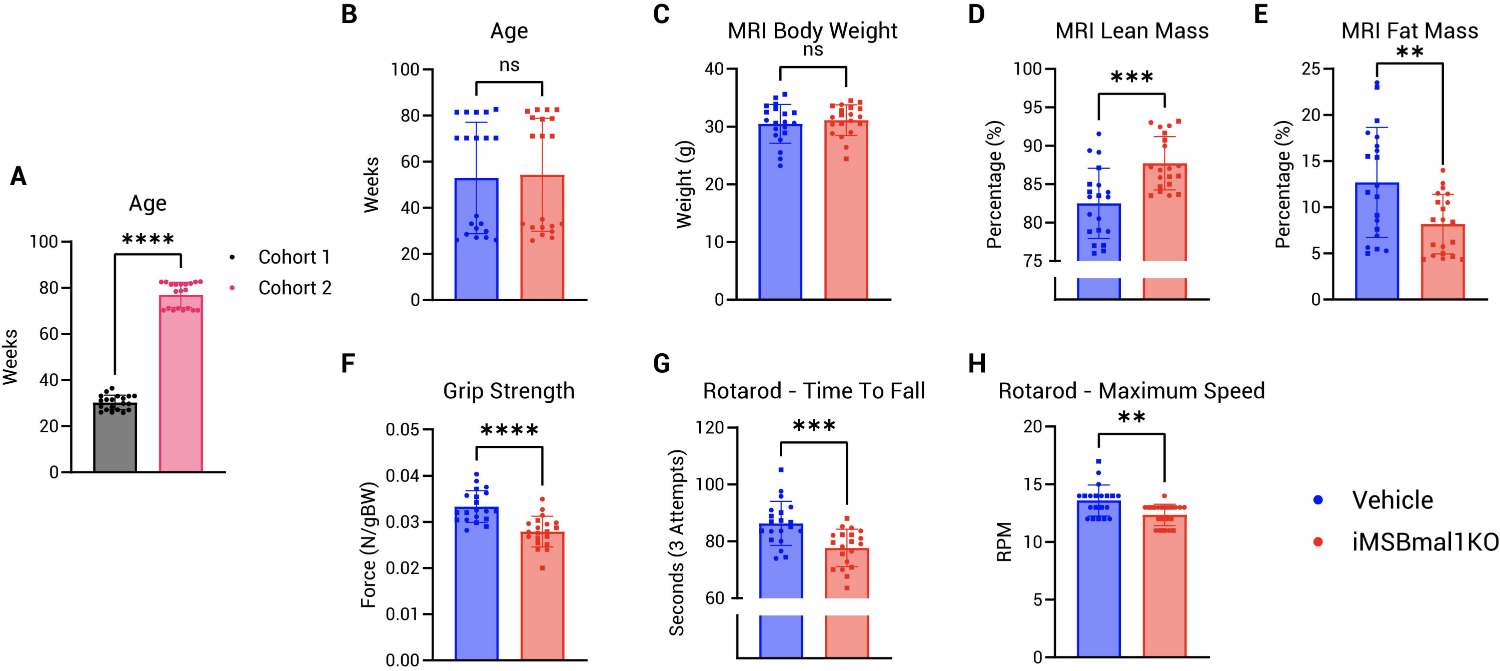

A
Age
****
Weeks
100
80
60
40
20
0
Cohort 1
Cohort 2
B
Age
ns
Weeks
C
MRI Body Weight
ns
Weight (g)
D
MRI Lean Mass
***
Percentage (%)
E
MRI Fat Mass
**
Percentage (%)
F
Grip Strength
****
Force (N/gBW)
G
Rotarod - Time To Fall
***
Seconds (3 Attempts)
H
Rotarod - Maximum Speed
**
RPM
Vehicle
iMSBmal1KO

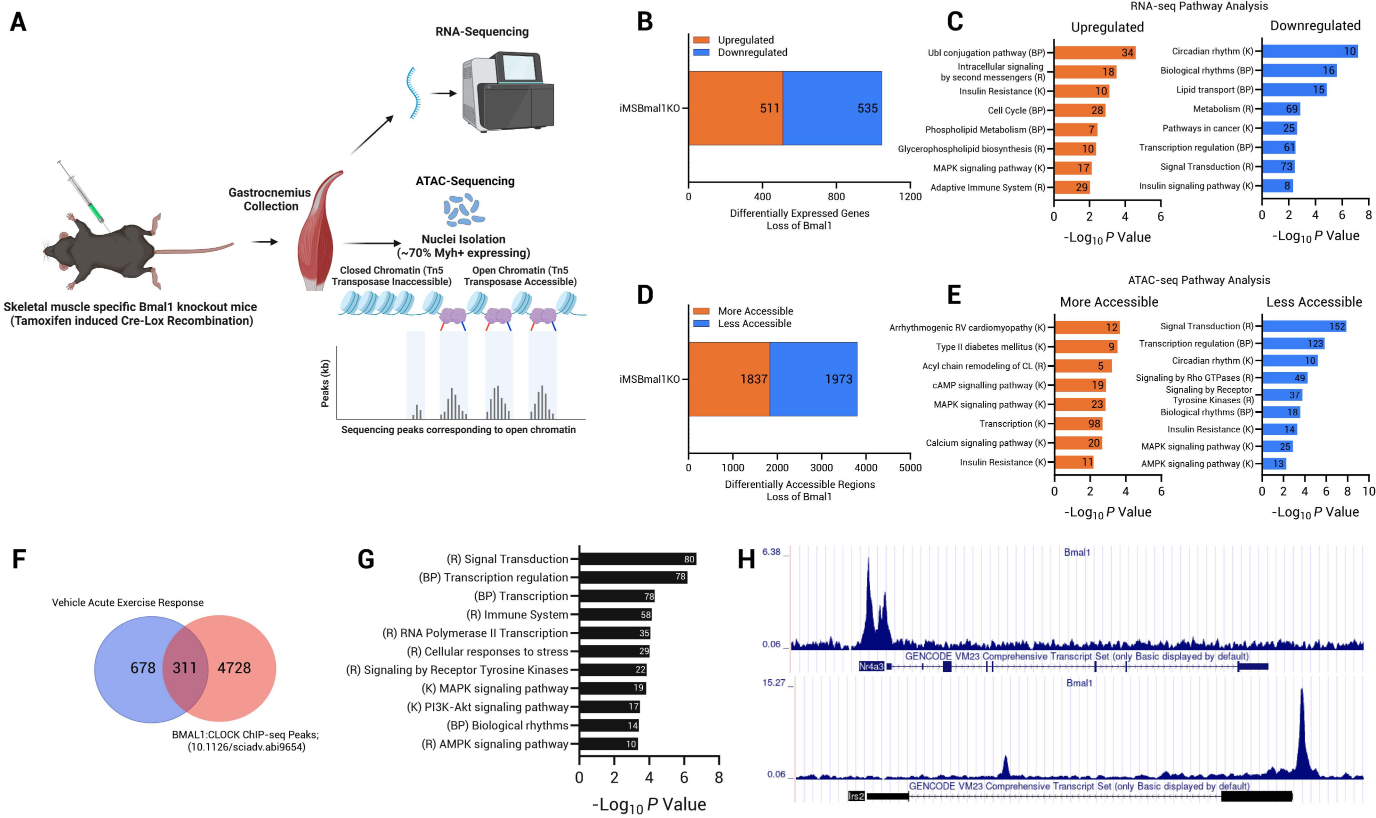
A
RNA-Sequencing
ATAC-Sequencing
Gastrocnemius Collection
Nuclei Isolation (~70% Myh+ expressing)
Closed Chromatin (Tn5 Transposase Inaccessible)
Open Chromatin (Tn5 Transposase Accessible)
Peaks (kb)
Sequencing peaks corresponding to open chromatin
Skeletal muscle specific Bmal1 knockout mice (Tamoxifen induced Cre-Lox Recombination)
B
Upregulated
Downregulated
iMSBmal1KO
511
535
0 400 800 1200
Differentially Expressed Genes Loss of Bmal1
C
RNA-seq Pathway Analysis
Upregulated
Ubl conjugation pathway (BP) 34
Intracellular signaling by second messengers (R) 18
Insulin Resistance (K) 10
Cell Cycle (BP) 28
Phospholipid Metabolism (BP) 7
Glycerophospholipid biosynthesis (R) 10
MAPK signaling pathway (K) 17
Adaptive Immune System (R) 29
Downregulated
Circadian rhythm (K) 10
Biological rhythms (BP) 16
Lipid transport (BP) 15
Metabolism (R) 69
Pathways in cancer (K) 25
Transcription regulation (BP) 61
Signal Transduction (R) 73
Insulin signaling pathway (K) 8
-Log10 P Value
D
More Accessible
Less Accessible
iMSBmal1KO
1837
1973
0 1000 2000 3000 4000 5000
Differentially Accessible Regions Loss of Bmal1
E
ATAC-seq Pathway Analysis
More Accessible
Arrhythmogenic RV cardiomyopathy (K) 12
Type II diabetes mellitus (K) 9
Acyl chain remodeling of CL (R) 5
cAMP signalling pathway (K) 19
MAPK signaling pathway (K) 23
Transcription (K) 98
Calcium signaling pathway (K) 20
Insulin Resistance (K) 11
Less Accessible
Signal Transduction (R) 152
Transcription regulation (BP) 123
Circadian rhythm (K) 10
Signaling by Rho GTPases (R) 49
Signaling by Receptor Tyrosine Kinases (R) 37
Biological rhythms (BP) 18
Insulin Resistance (K) 14
MAPK signaling pathway (K) 25
AMPK signaling pathway (K) 13
-Log10 P Value
F
Vehicle Acute Exercise Response
678
311
4728
BMAL1:CLOCK ChIP-seq Peaks; (10.1126/sciadv.abi9654)
G
(R) Signal Transduction 80
(BP) Transcription regulation 78
(BP) Transcription 78
(R) Immune System 58
(R) RNA Polymerase II Transcription 35
(R) Cellular responses to stress 29
(R) Signaling by Receptor Tyrosine Kinases 22
(K) MAPK signaling pathway 19
(K) PI3K-Akt signaling pathway 17
(BP) Biological rhythms 14
(R) AMPK signaling pathway 10
0 2 4 6 8
-Log10 P Value
H
6.38
Bmal1
0.06
GENCODE VM23 Comprehensive Transcript Set (only Basic displayed by default)
Nr4a3
15.27
Bmal1
0.06
GENCODE VM23 Comprehensive Transcript Set (only Basic displayed by default)
Irs2

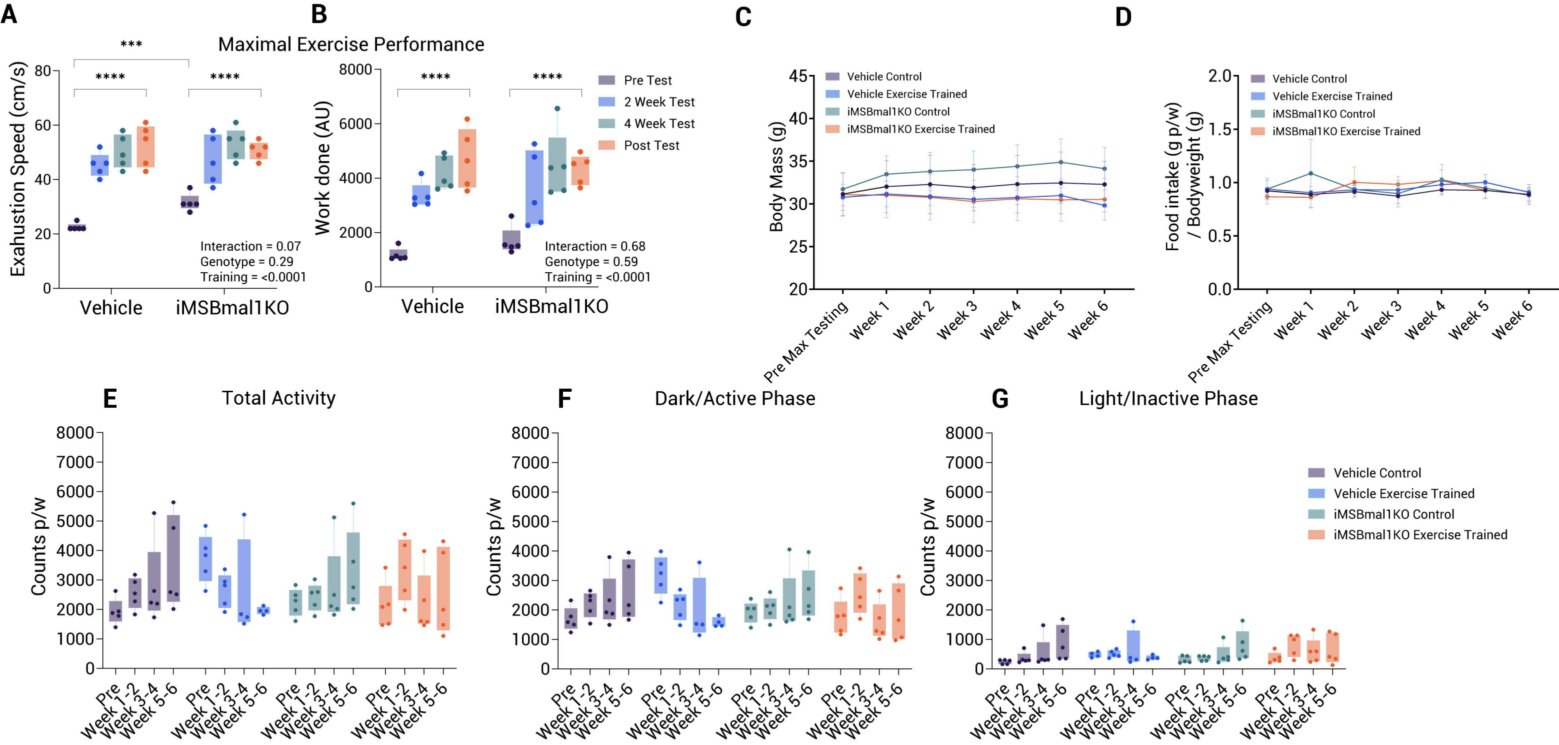
A
B
Maximal Exercise Performance
Exahustion Speed (cm/s)
Work done (AU)
***
****
****
Vehicle
iMSBmal1KO
Interaction = 0.07
Genotype = 0.29
Training = <0.0001
Interaction = 0.68
Genotype = 0.59
Training = <0.0001
Pre Test
2 Week Test
4 Week Test
Post Test
C
D
Body Mass (g)
Food intake (g p/w) / Bodyweight (g)
Vehicle Control
Vehicle Exercise Trained
iMSBmal1KO Control
iMSBmal1KO Exercise Trained
Pre Max Testing
Week 1
Week 2
Week 3
Week 4
Week 5
Week 6
E
Total Activity
F
Dark/Active Phase
G
Light/Inactive Phase
Counts p/w
Pre
Week 1-2
Week 3-4
Week 5-6

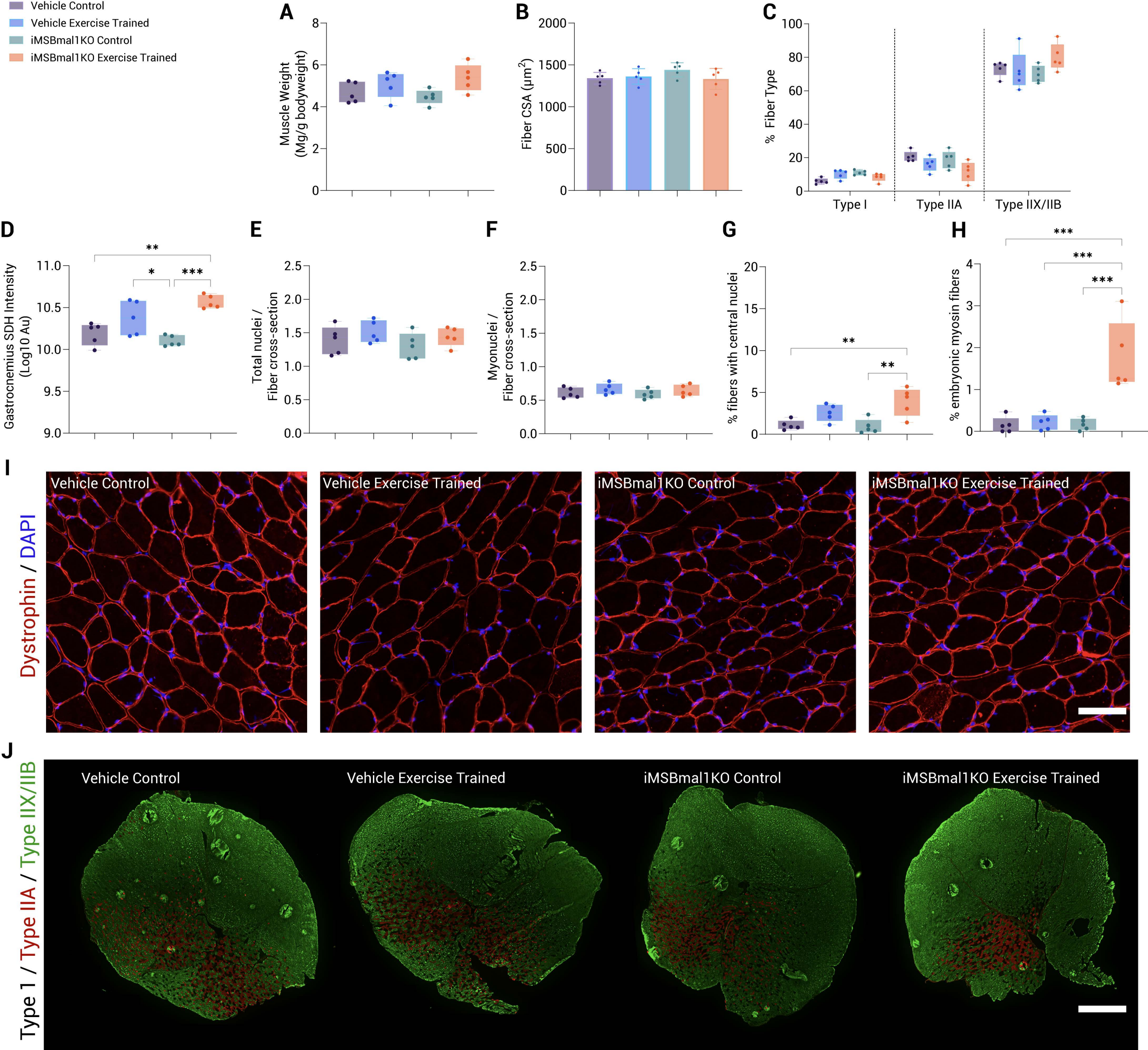
Vehicle Control
Vehicle Exercise Trained
iMSBmal1KO Control
iMSBmal1KO Exercise Trained
A
Muscle Weight (Mg/g bodyweight)
B
Fiber CSA (μm²)
C
% Fiber Type
Type I
Type IIA
Type IIX/IIB
D
Gastrocnemius SDH Intensity (Log10 Au)
E
Total nuclei / Fiber cross-section
F
Myonuclei / Fiber cross-section
G
% fibers with central nuclei
H
% embryonic myosin fibers
I
Dystrophin / DAPI
Vehicle Control
Vehicle Exercise Trained
iMSBmal1KO Control
iMSBmal1KO Exercise Trained
J
Type 1 / Type IIA / Type IIX/IIB
Vehicle Control
Vehicle Exercise Trained
iMSBmal1KO Control
iMSBmal1KO Exercise Trained

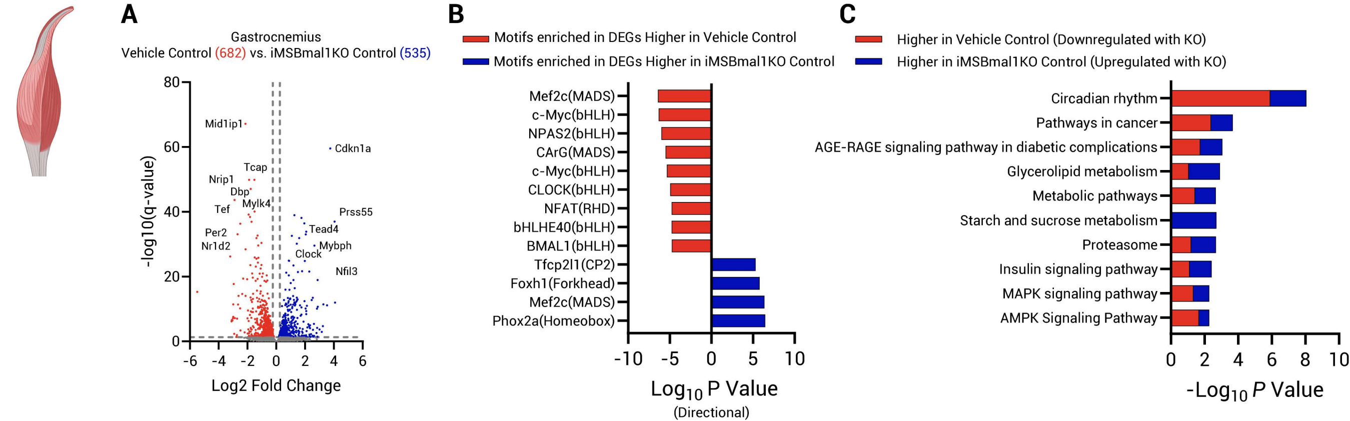
A
Gastrocnemius
Vehicle Control (682) vs. iMSBmal1KO Control (535)
-log10(q-value)
Log2 Fold Change
Mid1ip1
Cdkn1a
Tcap
Nrip1
Dbp
Mylk4
Tef
Prss55
Tead4
Per2
Nr1d2
Mybph
Clock
Nfil3
B
Motifs enriched in DEGs Higher in Vehicle Control
Motifs enriched in DEGs Higher in iMSBmal1KO Control
Mef2c(MADS)
c-Myc(bHLH)
NPAS2(bHLH)
CArG(MADS)
c-Myc(bHLH)
CLOCK(bHLH)
NFAT(RHD)
bHLHE40(bHLH)
BMAL1(bHLH)
Tfcp2l1(CP2)
Foxh1(Forkhead)
Mef2c(MADS)
Phox2a(Homeobox)
Log10 P Value
(Directional)
C
Higher in Vehicle Control (Downregulated with KO)
Higher in iMSBmal1KO Control (Upregulated with KO)
Circadian rhythm
Pathways in cancer
AGE-RAGE signaling pathway in diabetic complications
Glycerolipid metabolism
Metabolic pathways
Starch and sucrose metabolism
Proteasome
Insulin signaling pathway
MAPK signaling pathway
AMPK Signaling Pathway
-Log10 P Value

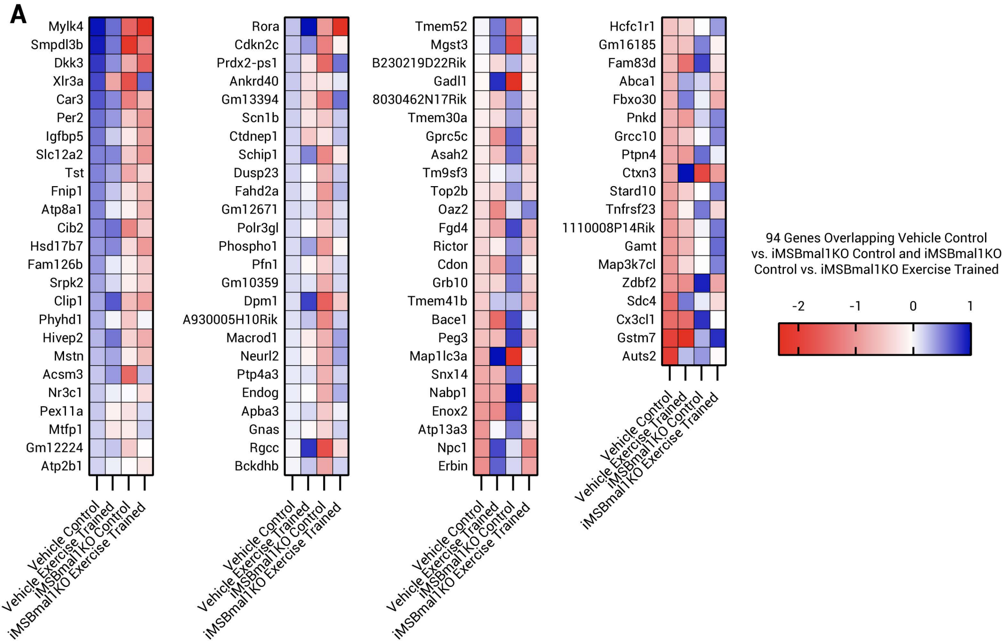
A
Mylk4
Smpdl3b
Dkk3
Xlr3a
Car3
Per2
Igfbp5
Slc12a2
Tst
Fnip1
Atp8a1
Cib2
Hsd17b7
Fam126b
Srpk2
Clip1
Phyhd1
Hivep2
Mstn
Acsm3
Nr3c1
Pex11a
Mtfp1
Gm12224
Atp2b1
Rora
Cdkn2c
Prdx2-ps1
Ankrd40
Gm13394
Scn1b
Ctdnep1
Schip1
Dusp23
Fahd2a
Gm12671
Polr3gl
Phospho1
Pfn1
Gm10359
Dpm1
A930005H10Rik
Macrod1
Neurl2
Ptp4a3
Endog
Apba3
Gnas
Rgcc
Bckdhb
Tmem52
Mgst3
B230219D22Rik
Gadl1
8030462N17Rik
Tmem30a
Gprc5c
Asah2
Tm9sf3
Top2b
Oaz2
Fgd4
Rictor
Cdon
Grb10
Tmem41b
Bace1
Peg3
Map1lc3a
Snx14
Nabp1
Enox2
Atp13a3
Npc1
Erbin
Hcfc1r1
Gm16185
Fam83d
Abca1
Fbxo30
Pnkd
Grcc10
Ptpn4
Ctxn3
Stard10
Tnfrsf23
1110008P14Rik
Gamt
Map3k7cl
Zdbf2
Sdc4
Cx3cl1
Gstm7
Auts2
Vehicle Control
Vehicle Exercise Trained
iMSBmal1KO Control
iMSBmal1KO Exercise Trained
94 Genes Overlapping Vehicle Control vs. iMSBmal1KO Control and iMSBmal1KO Control vs. iMSBmal1KO Exercise Trained
-2
-1
0
1

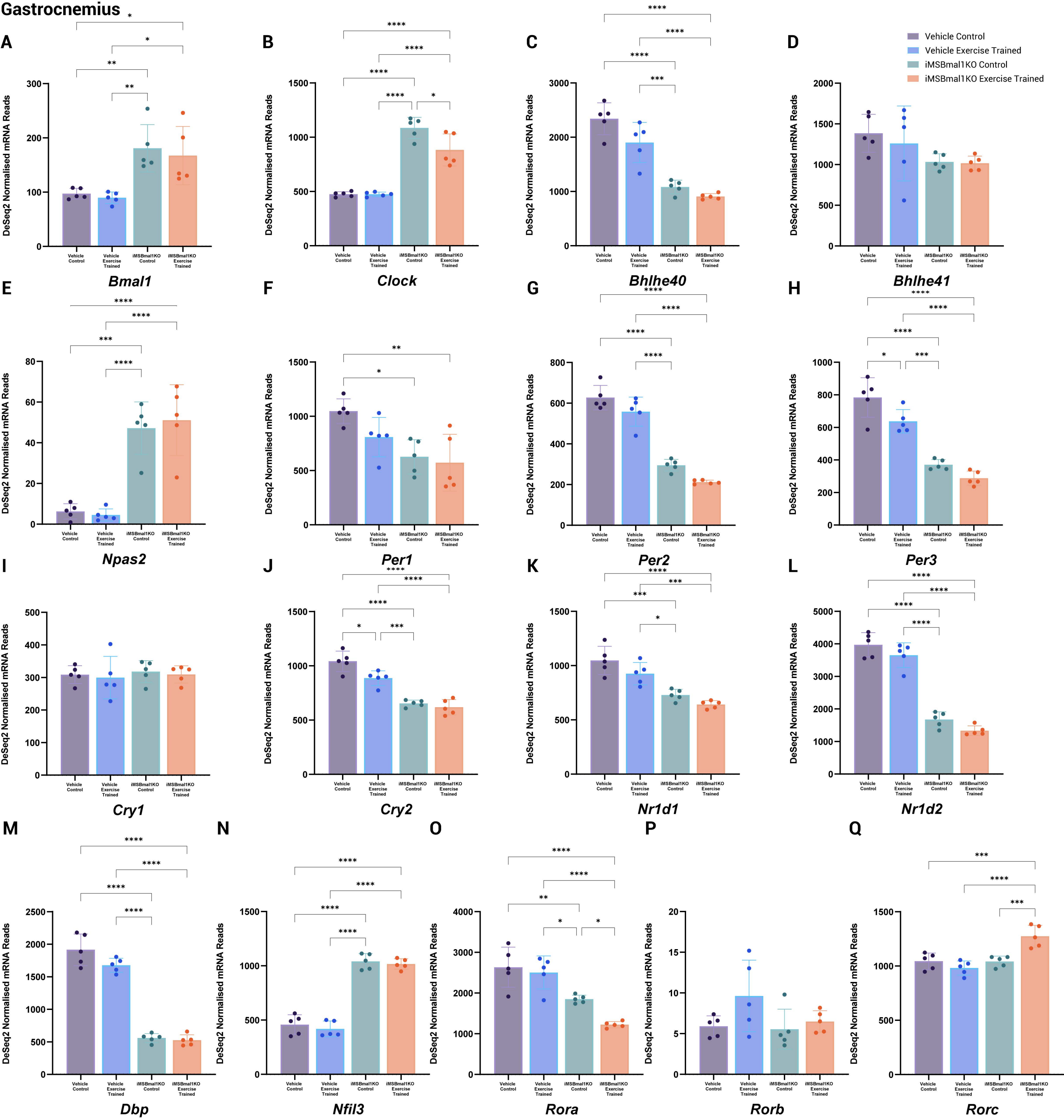
Gastrocnemius
A
B
C
D
Vehicle Control
Vehicle Exercise Trained
iMSBmal1KO Control
iMSBmal1KO Exercise Trained
DeSeq2 Normalised mRNA Reads
Bmal1
Clock
Bhlhe40
Bhlhe41
E
F
G
H
Npas2
Per1
Per2
Per3
I
J
K
L
Cry1
Cry2
Nr1d1
Nr1d2
M
N
O
P
Q
Dbp
Nfil3
Rora
Rorb
Rorc

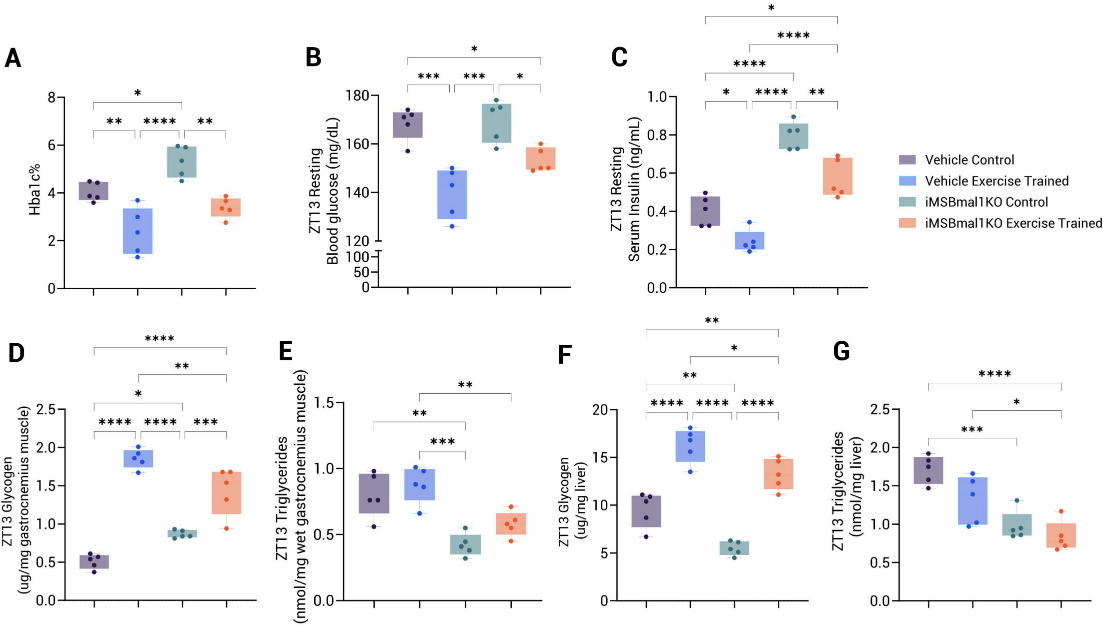

A
B
C
D
E
F
G
Hba1c%
ZT13 Resting Blood glucose (mg/dL)
ZT13 Resting Serum Insulin (ng/mL)
ZT13 Glycogen (ug/mg gastrocnemius muscle)
ZT13 Triglycerides (nmol/mg wet gastrocnemius muscle)
ZT13 Glycogen (ug/mg liver)
ZT13 Triglycerides (nmol/mg liver)
Vehicle Control
Vehicle Exercise Trained
iMSBmal1KO Control
iMSBmal1KO Exercise Trained

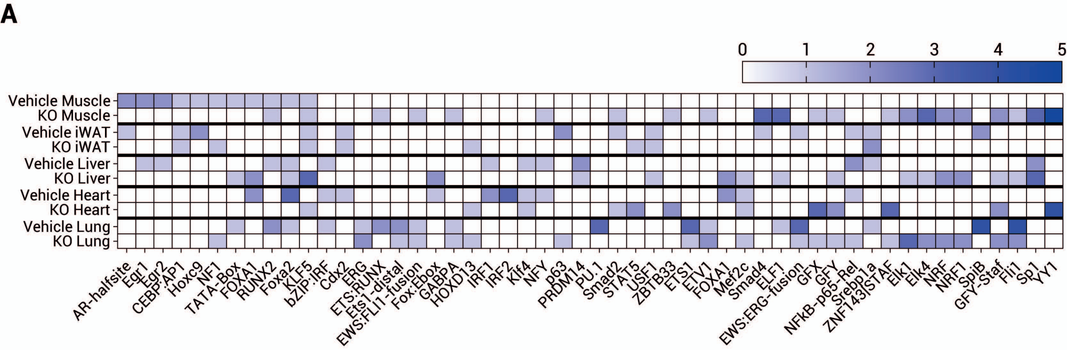
A
0
1
2
3
4
5
Vehicle Muscle
KO Muscle
Vehicle iWAT
KO iWAT
Vehicle Liver
KO Liver
Vehicle Heart
KO Heart
Vehicle Lung
KO Lung
AR-halfsite
Egr1
Egr2
CEBP:AP1
Hoxc9
NF1
TATA-Box
FOXA1
RUNX2
Foxa2
KLF5
bZIP:IRF
Cdx2
ERG
ETS:RUNX
Ets1-distal
EWS:FLI1-fusion
Fox:Ebox
GABPA
HOXD13
IRF1
IRF2
Klf4
NFY
p63
PRDM14
PU.1
Smad2
STAT5
USF1
ZBTB33
ETS1
ETV1
FOXA1
Mef2c
Smad4
ELF1
EWS:ERG-fusion
GFX
GFY
NFkB-p65-Rel
Srebp1a
ZNF143|STAF
Elk1
Elk4
NRF
NRF1
SpiB
GFY-Staf
Fli1
Sp1
YY1

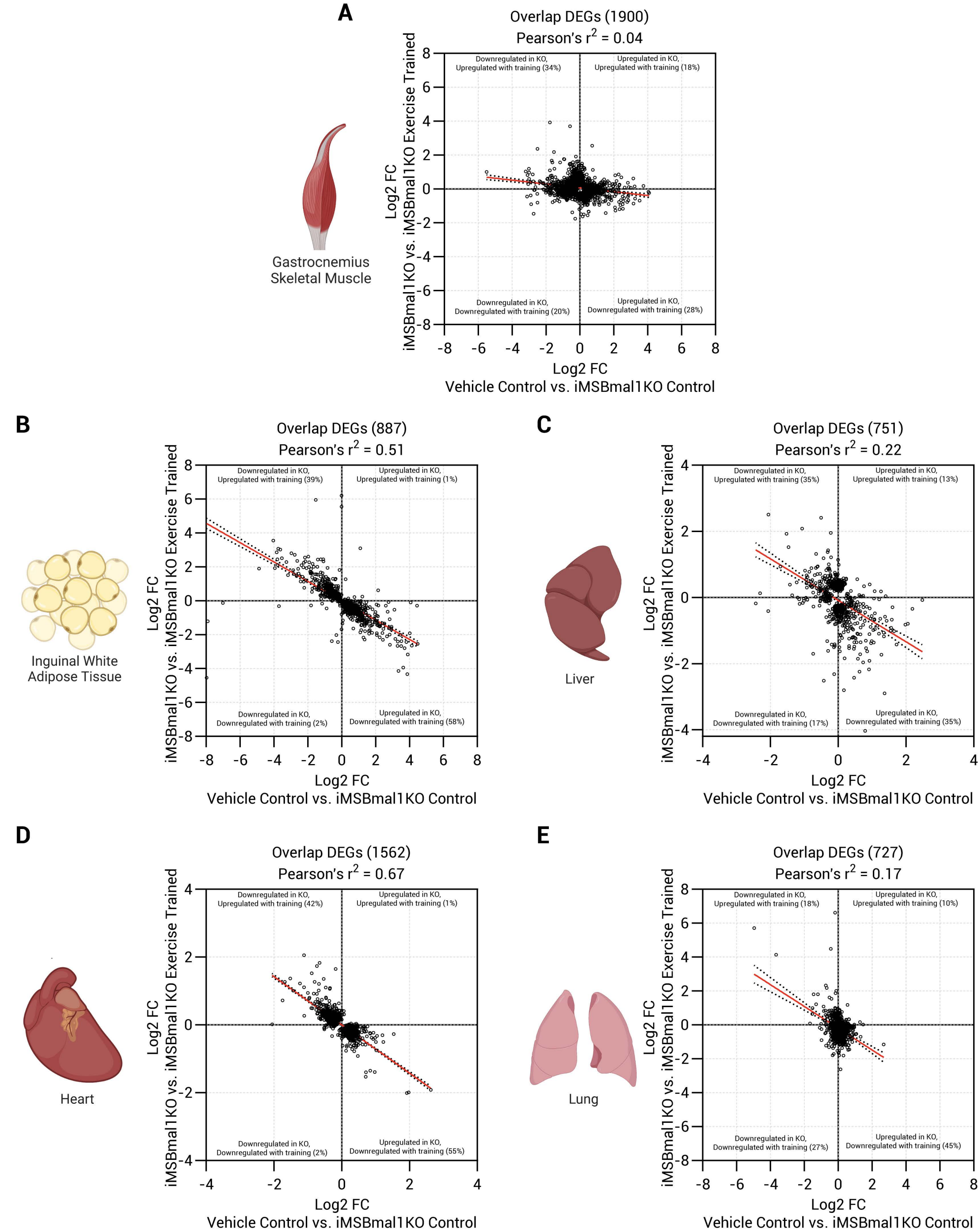
A
Overlap DEGs (1900)
Pearson's r2 = 0.04
Gastrocnemius Skeletal Muscle
Downregulated in KO, Upregulated with training (34%)
Upregulated in KO, Upregulated with training (18%)
Downregulated in KO, Downregulated with training (20%)
Upregulated in KO, Downregulated with training (28%)
iMSBmal1KO vs. iMSBmal1KO Exercise Trained Log2 FC
Log2 FC Vehicle Control vs. iMSBmal1KO Control
B
Overlap DEGs (887)
Pearson's r2 = 0.51
Inguinal White Adipose Tissue
Downregulated in KO, Upregulated with training (39%)
Upregulated in KO, Upregulated with training (1%)
Downregulated in KO, Downregulated with training (2%)
Upregulated in KO, Downregulated with training (58%)
C
Overlap DEGs (751)
Pearson's r2 = 0.22
Liver
Downregulated in KO, Upregulated with training (35%)
Upregulated in KO, Upregulated with training (13%)
Downregulated in KO, Downregulated with training (17%)
Upregulated in KO, Downregulated with training (35%)
D
Overlap DEGs (1562)
Pearson's r2 = 0.67
Heart
Downregulated in KO, Upregulated with training (42%)
Upregulated in KO, Upregulated with training (1%)
Downregulated in KO, Downregulated with training (2%)
Upregulated in KO, Downregulated with training (55%)
E
Overlap DEGs (727)
Pearson's r2 = 0.17
Lung
Downregulated in KO, Upregulated with training (18%)
Upregulated in KO, Upregulated with training (10%)
Downregulated in KO, Downregulated with training (27%)
Upregulated in KO, Downregulated with training (45%)
